# GLUT1 is redundant in hypoxic and glycolytic nucleus pulposus cells of the intervertebral disc

**DOI:** 10.1101/2022.07.22.501129

**Authors:** Shira N. Johnston, Elizabeth S. Silagi, Vedavathi Madhu, Duc H. Nguyen, Irving M. Shapiro, Makarand V. Risbud

## Abstract

Glycolysis is central to homeostasis of nucleus pulposus (NP) cells in the avascular intervertebral disc. Since the glucose importer, GLUT1, is a highly enriched phenotypic marker of NP cells, we hypothesized that it is vital for the development and post-natal maintenance of the disc. Surprisingly, primary NP cells treated with two well-characterized GLUT1 inhibitors maintained normal rates of glycolysis and ATP production, indicating intrinsic compensatory mechanisms. We show in vitro that NP cells mitigate GLUT1 loss by rewiring glucose import through GLUT3. Noteworthy, we demonstrate that substrates, such as glutamine and palmitate, do not compensate for glucose restriction resulting from dual inhibition of GLUT1/3 and inhibition compromises long-term cell viability. To investigate the redundancy of GLUT1 function in NP, we generated two NP-specific knockout mice: Krt19^CreERT^; Glut1^f/f^ and Foxa2^Cre^; Glut1^f/f^. Noteworthy, there were no apparent defects in post-natal disc health or development and maturation in mutant mice. Microarray analysis confirmed that GLUT1 loss did not cause transcriptomic alterations in the NP, supporting that cells are refractory to GLUT1 loss. These observations provide the first evidence of functional redundancy in GLUT transporters in the physiologically hypoxic intervertebral disc and underscore the importance of glucose as the indispensable substrate for NP cells.

## INTRODUCTION

The phenotype of nucleus pulposus (NP) cells reflects their notochordal origin and unique hypoxic environment (1–3). Indeed, an initial definition of the NP cell phenotype was based on the observation that the HIF-1α transcription factor and glucose transporter, GLUT1, were highly enriched in the NP and not the annulus fibrosus (AF) compartment of the intervertebral disc (4). To facilitate hypoxic adaptation, NP cells constitutively express HIF-1α, given that the vasculature in the adjacent vertebral bodies does not penetrate the NP compartment (5, 6). HIF-1α promotes the biosynthetic activity of NP cells by regulating glycolytic metabolism and mitochondrial TCA cycle flux and conditional deletion of HIF-1α in notochord results in massive NP cell apoptosis at birth likely due to metabolic failure (7–9). Further investigations into the HIF-1α transcriptional program showed that Glut1 was the HIF-1 target and expression was regulated through PHD3-dependent modulation of HIF-1α-C-TAD activity (10).

It has been well established that the maintenance of glycolytic flux and nutrient-metabolite balance is critical for cell survival in the intervertebral disc. Glucose passively diffuses from the vertebral capillaries, through the hyaline cartilaginous endplates and proteoglycan-rich extracellular matrix of the NP tissue compartment to reach the resident NP cells at the center of the disc. Despite this long diffusion distance, glucose levels must surpass a critical threshold for cells to remain viable; glucose concentrations below 0.5 mM are shown to promote cell death (11). Numerous studies have confirmed that glucose availability is required for critical NP cellular processes such as protein and proteoglycan biosynthesis, glycolytic flux, and maintaining cell viability (12–15). Furthermore, disruption in the balance between glucose consumption and lactic acid production can significantly impact NP cell physiology (16, 17). Several factors can influence the rate of glucose consumption in NP cells, including nutrient deprivation from decreased glucose diffusivity or increased cell density (18); pH buffering capacity governed by relative levels of bicarbonate and sodium (19); mechanical stress (20), and oxygen tension (21, 22).

Despite the well-studied importance of glucose availability and consumption on NP cell viability *in vitro*, few studies have focused on the relationship between glucose consumption and disc health *in vivo*. This is due to the complexity of studying solute transport and metabolite concentrations in animal models or by using genetic techniques. However, some studies have shown enriched expression of GLUT1 in human NP and that GLUT3 and GLUT9 levels were lower than GLUT1 (23). These results suggest that glucose transporter redundancy may be required for disc health and function.

Considering that GLUT1 is a high affinity glucose transporter with nearly ubiquitous expression in all tissue types, it is not surprising that embryos with homozygous GLUT1 deficiency are nonviable (24). GLUT1 haploinsufficiency in mice causes profound developmental defects recapitulating those seen in human patients with GLUT1 deficiency syndrome (25). Both humans and mice with GLUT1 haploinsufficiency experience microcephaly, impaired motor function, epileptiform changes on EEG, and hypoglycorrhachia (25). Due to the severe impact of loss of GLUT1 on gross embryonic development, tissue-specific knock-out mice have been used to delineate the role of GLUT1 in developing skeleton and other connective tissues (26–30). These studies show that GLUT1 deletion leads to metabolic reprogramming of cells, profound phenotypic changes, and compromised tissue function. In skeletal tissues, loss of GLUT1 expression results in severely impaired bone development, emphasizing the importance of GLUT1 in the maintenance of bone health. As far as mechanisms are concerned, it is shown that GLUT1 expression precedes Runx2, and is required for promoting bone formation by blocking AMPK-dependent degradation of Runx2 (27). Mouse models of *Glut1* loss of function in the growth plate and articular cartilage demonstrate that *Glut1* function is required for cartilage structure and function regulating cell proliferation, matrix production, and resistance to injury and osteoarthritis (31). GLUT1 expression is controlled by a unique BMP-mTORC1-HIF1 signaling cascade in chondrocytes where it is required for chondrocyte proliferation and hypertrophy (28).

Since GLUT1-mediated glycolytic metabolism plays a fundamental role in many tissues including bone and cartilage homeostasis, we surmised that loss of GLUT1 expression in the NP would impact both development and age-related structure of the disc (26–30). However, unlike the other skeletal tissues, both conditional and inducible loss of GLUT1 expression in the NP did not result in notable degenerative changes in the discs of developing perinatal mice or in skeletally mature mice. Surprisingly, microarray analysis of global transcriptomic changes in NP tissue isolated from conditional GLUT1 knock-out mice did not reveal any differentially regulated genes besides *Slc2a1* (encoding GLUT1) – a finding which suggests NP cells are refractory to loss of GLUT1. In fact, long-term GLUT1 inhibition had no effect on the rates of NP glycolytic flux or oxidative metabolism; these findings indicate that NP cells can potentially mitigate the loss of GLUT1 function by rewiring glucose import through GLUT3. Importantly, our findings suggest that under glucose limiting conditions resulting from dual inhibition of GLUT1/3, NP cells do not evidence metabolic reprogramming to utilize alternative substrates, such as glutamine and fatty acids. These findings provide the first evidence of functional redundancy in GLUT transporters in a physiologically hypoxic intervertebral disc and underscore the importance of glucose as the indispensable metabolic substrate for NP cells.

## RESULTS

### HIF-1 dependent GLUT1 expression is highly enriched in the NP and declines with age

During embryogenesis, the notochord and developing nucleus pulposus (NP) compartment of the intervertebral disc is hypoxic and exhibits robust HIF-1α activity. Our previous work has shown that notochord-specific HIF-1α deletion in Foxa2^Cre^; HIF-1α^f/f^ (HIF-1α^cKO^) mice leads to massive NP cell death at birth, likely due to metabolic failure of cells that rely primarily on glycolytic metabolism for their energetic needs (7–9). It is therefore not surprising that through early development to skeletal maturity expression of the glucose transporter, *Slc2a1* (GLUT1) is highly enriched in NP cells (Figure 1A) and is regarded as a phenotypic marker (3, 32, 33). In fact, GLUT1 expression is substantially decreased in the NP of E15.5 HIF-1α^cKO^ mice without affecting level of carbonic anhydrase 3 (CA3), another hypoxia-sensitive NP-phenotypic marker (34) (Figure 1, B and B’). These results indicate that loss of GLUT1 and consequent restriction on glucose availability shortly precedes the catastrophic NP cell death observed in HIF-1α^cKO^ mice at birth. Furthermore, changes in GLUT1 expression in the intervertebral disc correlate to both age and degenerative state (23). Our studies show that in mice, GLUT1 expression is significantly decreased during normal aging from 1 month to 24 months (Figure 1, C-D). Immunofluorescence staining of GLUT1 shows a robust expression of GLUT1 in the NP compartment at early time points, followed by a significant decrease in GLUT1 abundance by 24 months of age (Figure 1, C and C’). In fact, Western blot confirmed that GLUT1 protein level was markedly decreased by as early as 14 months of age (Figure 1D). Based on these findings we hypothesized that GLUT1 is critically important for NP cell survival and function.

**Figure 1.**
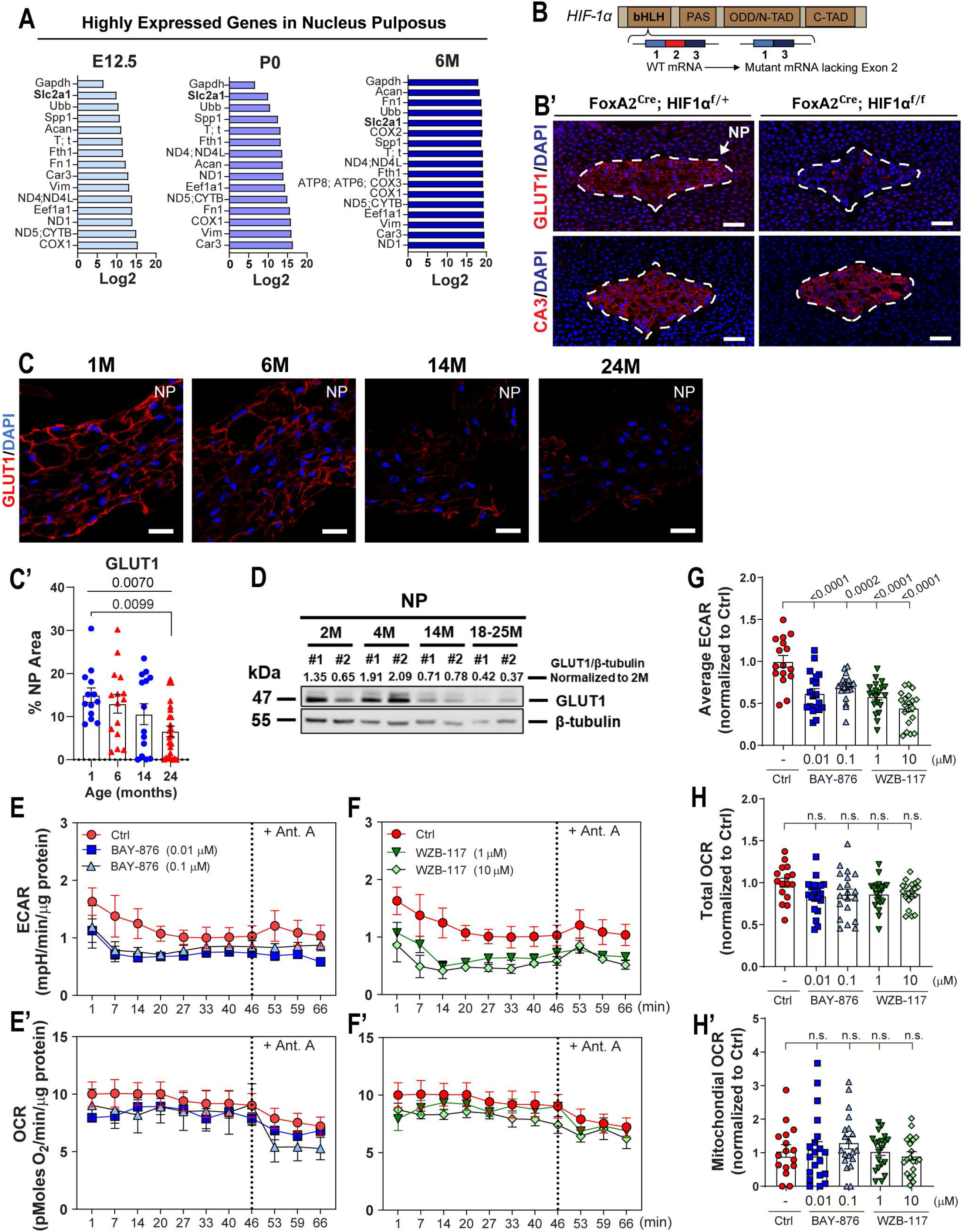
GLUT1, an enriched NP marker is a HIF-1 target and its expression declines with aging. (A) Graph showing some of the most highly expressed in mouse NP cells at E12.5, P0 and 6-months (GSEs: GSE100934 and GSE134955). (B) Schematic showing deletion of HIF-1α exon 2 to generate mutant HIF-1α mRNA. (B’) Representative IHC images of NP cell phenotypic markers GLUT1 and CA3 in HIF-1α WT and HIF-1α^cKO^ (Foxa2^Cre^; HIF-1α^f/f^) mice (scale bar = 50 μm). (C, C’) Representative images and quantification of GLUT1 in C57BL6/J mice with aging (scale bar = 25 μm) (n = 5 mice/timepoint; 3-6 discs/animal; 15-30 discs/timepoint). (D) Western blot showing GLUT1 levels in mouse NP with aging (n = 2 mice/timepoint; 20 discs/animal were pooled). (E, E’) Effect of short-term (1 hour) GLUT1 inhibition by BAY-876 on ECAR and OCR measurements in NP cells. (F, F’) Effect of short-term GLUT1 inhibition by WZB-117 on ECAR and OCR measurements in NP cells. (G) Short-term GLUT1 inhibition by BAY-876 and WZB-117 results in decreased average ECAR. (H) Short-term GLUT1 inhibition does not affect total OCR in BAY-876 and WZB-117 treated cells. (H’) Mitochondrial OCR measurements following short term treatment by BAY-876 and WZB-117. (n = 6 independent experiments, 4 technical replicates/experiment/group). Quantitative measurements represent mean ± SEM. Significance was determined using a one-way ANOVA (G-H’) or Kruskal Wallis (C’) with Dunnett’s or Dunn’s post hoc test as appropriate.

### Long-term inhibition of GLUT1 does not affect glycolytic or oxidative metabolism in NP cells

To determine if the loss of GLUT1 function directly impairs NP cell metabolism, we modeled the loss of GLUT1 in NP cells *in vitro* with two highly specific pharmacological inhibitors; namely, BAY-876 and WZB-117 (35, 36). Using a Seahorse Flux Analyzer, we assessed metrics of NP cell glycolytic flux by measuring Extracellular Acidification Rate (ECAR) and oxidative flux by oxygen consumption rate (OCR). After short-term inhibition of GLUT1 for 1 hour, NP cells showed significantly decreased average ECAR with both inhibitors (Figure 1, E-G). However, total OCR and mitochondrial OCR remained unchanged, suggesting the NP cells did not undergo a metabolic switch from glycolytic to oxidative metabolism (Figure 1, E’, F’, H, and H’). Moreover, inhibition of the electron transport chain by antimycin A failed to influence mitochondrial respiration. It is concluded that while the baseline mitochondrial respiration remains low in both control and inhibitor-treated cells this is not related to failure to provide metabolic intermediates to power the electron transport chain (Figure 1, E’ and F’).

To determine if longer-term inhibition of GLUT1 influences NP cell metabolism, we measured ECAR, OCR, and calculated the ATP production rates from glycolysis and oxidative metabolism in NP cells treated with two concentrations of BAY-876 and WZB-117 for 24h. We recorded raw ECAR and OCR traces under basal conditions (no glucose), followed by sequential addition of glucose (substrate), oligomycin (ATP-synthase inhibitor), and rotenone + myxothiozol (ETC inhibitors) (Figure 2, A-B’). Surprisingly, there were no significant differences in the average ECAR and OCR between control and GLUT1-inhibited NP cells (Figure 2, C and D), as is also evident from the raw tracer profiles. Using the published method by Mookerjee and colleagues (37), we calculated glycolytic and oxidative ATP production rates following GLUT1 inhibition with BAY-876 and WZB-117. We estimate that under basal conditions (no glucose), oxidative metabolism generates ∼50% of ATP in control NP cells, however, this decreases to ∼10-25% when glucose is added (Figure 2, E and E’). Importantly, our results showed that there was no difference in ATP production rates from glycolysis or oxidative phosphorylation in GLUT1-inhibited cells as compared to controls (Figure 2, E and E’). Furthermore, blocking oxidative ATP production entirely with oligomycin showed strikingly little effect on glycolytic ATP production rate, suggesting that glycolytic flux is largely independent of oxidative metabolism in NP cells (Figure 2, E and E’). To understand if there was an effect of GLUT1 loss on glycolytic reserve, we measured ECAR, OCR, and calculated the proton production rate (PPR) in NP cells treated with two concentrations of BAY-876 and WZB-117 for 24h. We recorded ECAR and OCR traces under basal conditions (no glucose), followed by sequential addition of glucose, rotenone + myxothiozol (ETC inhibitors), and Carbonyl cyanide *p*-(tri-fluromethoxy)phenyl-hydrazone (FCCP) an uncoupler of mitochondrial oxidative phosphorylation + monensin (Mon) which decreases the mitochondrial membrane potential (Figure 2, F, F’, G, and G’). There were minimal differences in the average ECAR and OCR between control and GLUT1-inhibited NP cells (Figure 2, F, F’, G, and G’) and accordingly a minimal difference in the PPR and glycolytic reserve of GLUT1-inhibited cells as compared to controls (Figure 2, F’’ and G’’). Taken together, the results of the Seahorse metabolic assays suggested that inhibiting glucose uptake though GLUT1 in NP cells causes an immediate decrease in glycolytic flux. However, compensatory mechanisms are capable of restoring NP cell glycolytic metabolism within 24 hours, without initiating a metabolic shift towards oxidative metabolism or affecting the glycolytic capacity and reserve of the cells.

**Figure 2.**
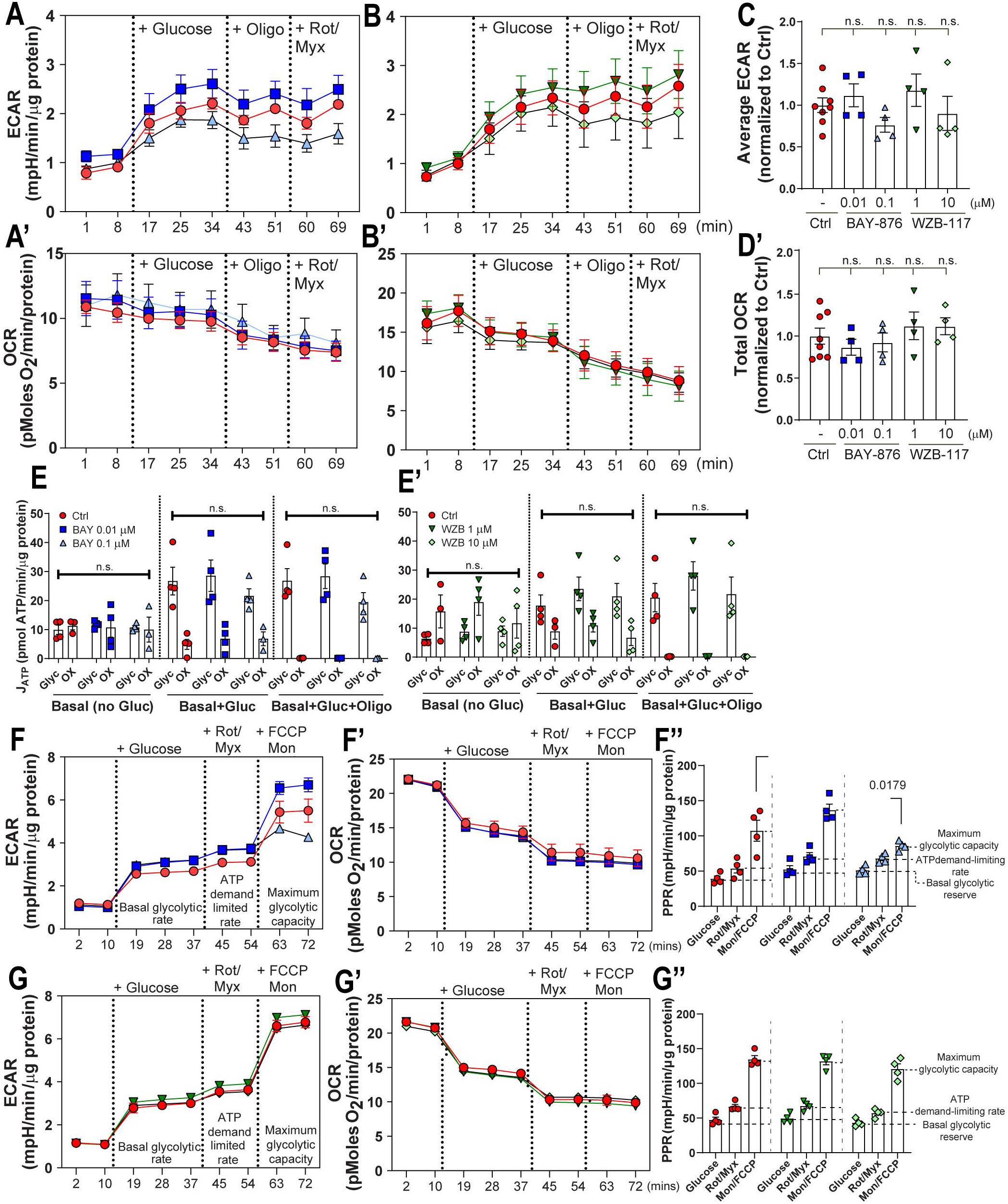
Long-term inhibition of GLUT1 does not affect glycolytic or oxidative metabolism in NP cells. (A-A’) ECAR and OCR profiles following long term GLUT1 inhibition (24 h) by BAY-876 in NP cells. (B-B’) ECAR and OCR profiles following long term GLUT1 inhibition (24 h) by WZB-117. (C-D) Long term GLUT1 inhibition by BAY-876 and WZB-117 did not alter (C) ECAR and (D) OCR measurements. (E-E’) Long term GLUT1 inhibition by (E) BAY-876 and (E’) WZB-117 did not affect ATP production by NP cells. (F-F’) ECAR and OCR traces in 24h BAY-876 treated and control NP cells to assess glycolytic capacity and reserve. (F’’) Inhibition of GLUT1 by BAY-876 minimally impacts glycolytic reserve by NP cells only at the higher concentration of 0.1 μM. (G-G’) ECAR and OCR traces in 24h WZB-117 treated and control NP cells to assess proton production rate. (G’’) Inhibition of GLUT1 by WZB-117 does not alter proton production rate by NP cells. Quantitative measurements represent mean ± SEM (n = 4 biological replicates, 4 technical replicates/experiment/group). Significance was determined using one-way ANOVA.

### GLUT3 sustains glucose uptake in absence of GLUT1

Although GLUT1 is the highest expressed transporter in the NP, other glucose transporters including hypoxia-sensitive GLUT3 and GLUT9 are reported to be expressed in the NP and may also facilitate glucose uptake (3, 23). To test this hypothesis, we evaluated if GLUT1 is solely required for glucose import by measuring glucose uptake in NP cells treated with GLUT1 inhibitors, BAY-876/WZB-117, and the potent GLUT1/2/3 inhibitor, Glutor. We treated primary NP cells with the glucose mimic, 2-deoxyglucose (2-DG), and measured the subsequent intracellular accumulation of 2-deoxyglucose-6-phosphate, which cannot undergo glycolysis. The 2-DG uptake assay clearly showed that simultaneously blocking GLUT1 and GLUT3 decreased glucose uptake by up to 80 to 90% compared to control in a dose and time-dependent manner (Figure 3, A and A’). Interestingly, GLUT1 inhibition alone for 6 and 24 hours with BAY-876 or WZB-117 did not result in decreased glucose uptake implying a rapid compensation for loss of GLUT1 function. These findings also suggest that alternative glucose transporters, such as GLUT3 (NP cells do not express GLUT2 which has a high Km for glucose of ∼17.1 mM) compensate for GLUT1 and may be responsible for much of the glucose uptake in NP cells. These results raised a possibility that simultaneous long-term inhibition of GLUT1 and GLUT3 will compromise NP cell survival. We, therefore, measured NP cell viability following Glutor treatment for up to 72 hours. It was evident that cell viability decreased with increasing Glutor dose and the time of treatment (Figure 3, B-B’’).

**Figure 3.**
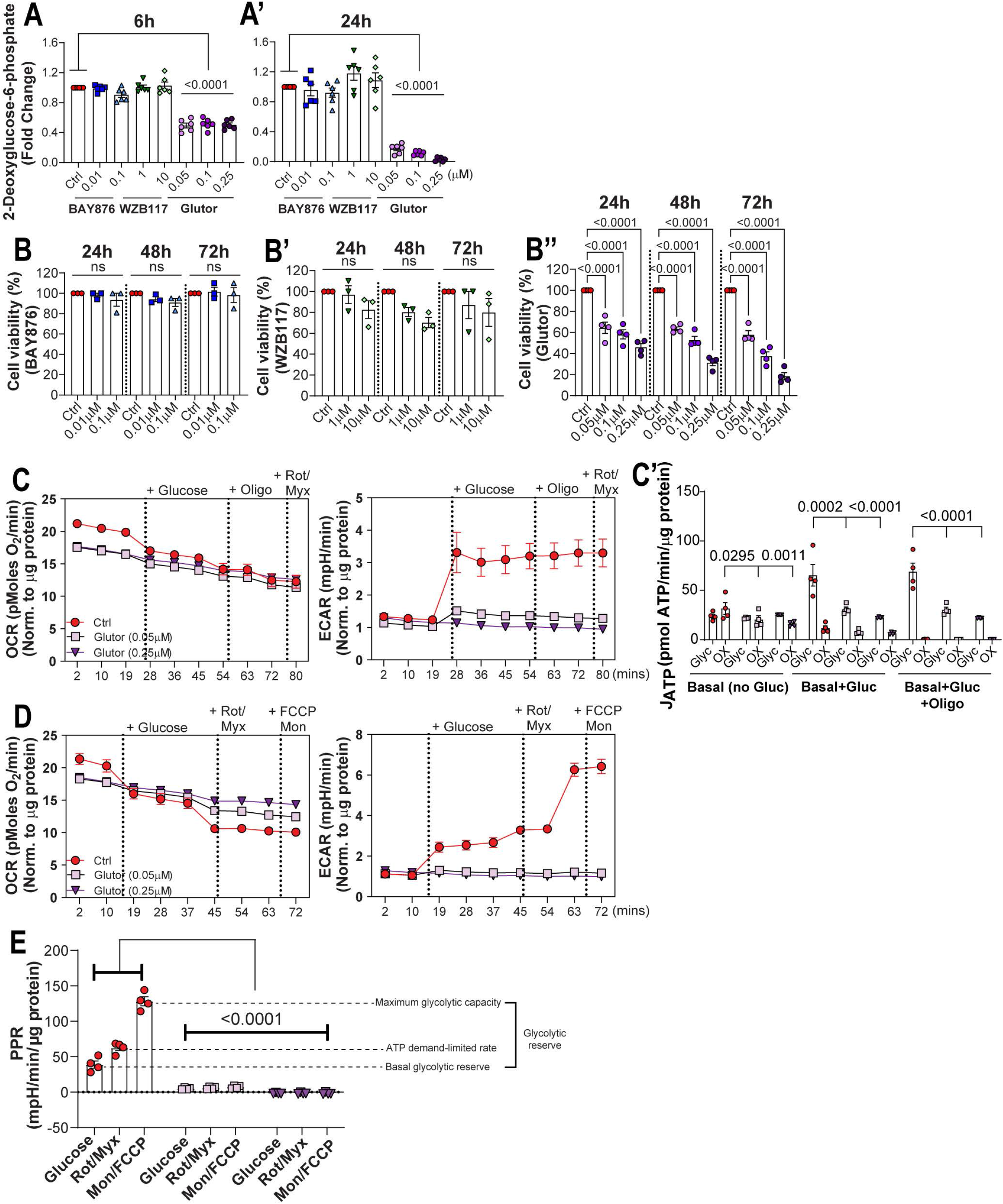
GLUT3 facilitates uptake of glucose, a metabolic substrate indispensable for NP cells. (A, A’) 2-Deoxyglucose (2-DG) uptake in cells treated with GLUT1 inhibitors and GLUT1/3 dual inhibitor Glutor from (A) 6 to (A’) 24 hours. (B-B”) Cell viability following GLUT1 (BAY-876 and WZB-117) and GLUT1/3 inhibition (Glutor). (C) OCR and ECAR traces in 24h Glutor treated and control NP cells to assess ATP production rate. (C’) Inhibition of GLUT1 and GLUT3 by Glutor significantly decreased the ATP production by NP cells. (D) OCR and ECAR traces in 24h Glutor treated and control NP cells to assess proton production rate. (E) Inhibition of GLUT1 and GLUT3 by Glutor significantly decreased the proton production rate in NP cells. Quantitative measurements represent mean ± SEM (n = 4-6 biological replicates, 4 technical replicates/experiment/group). Significance of differences was determined using one-way ANOVA with Sidak’s or Dunnett’s post hoc test as appropriate.

To ascertain if GLUT1/3 inhibition impacts NP cell metabolism, we performed seahorse assays to measure ATP production rate and glycolytic flux of NP cells treated with two concentrations of Glutor for 24h (Figure 3, C and D). Glutor-treated NP cells did not show an increase in average ECAR and a corresponding decrease in average OCR following glucose addition and co-treatment with mitochondrial inhibitors as evidenced by untreated controls (Figure 3, C and D). Importantly, there was a significant decrease in glycolytic ATP production rate in Glutor-treated cells in the presence of glucose with or without Oligomycin which inhibits mitochondrial ATP generation (Figure 3C’). Moreover, Glutor-treatment resulted in a complete and striking collapse of glycolytic capacity and reserve and compared to control cells (Figure 3E). These results clearly indicated that in NP cells, glucose uptake is sustained by both GLUT1 and GLUT3, and supports the notion that GLUT3 can compensate for GLUT1 loss.

Moreover, similar to GLUT1, there was a robust decrease in GLUT3 levels in the NP of E15.5 HIF-1α^cKO^ mice suggesting that pronounced cell death observed at birth in the NP compartment of these cKO mice could be in part due to constrained availability of glucose aggravating metabolic failure (9) (Figure 4A, Supplemental Figure 1A). Likewise, we observed a strong age-dependent decrease in GLUT3 abundance in the NP compartment with a significant reduction noted at 24 months (Supplemental Figure 1, B-D). Furthermore, qRT-PCR analysis showed a significant decrease in mRNA levels of *Glut3, Glut9*, and *Sglt1* with aging (Supplemental Figure 2A). Interestingly however, a significant increase in *Glut1*/*Slc2a1* mRNA was noted at 24 months suggesting a plausible compensation for the loss of GLUT1. Together these results suggest a correlation of glucose availability to age-dependent degeneration.

**Figure 4.**
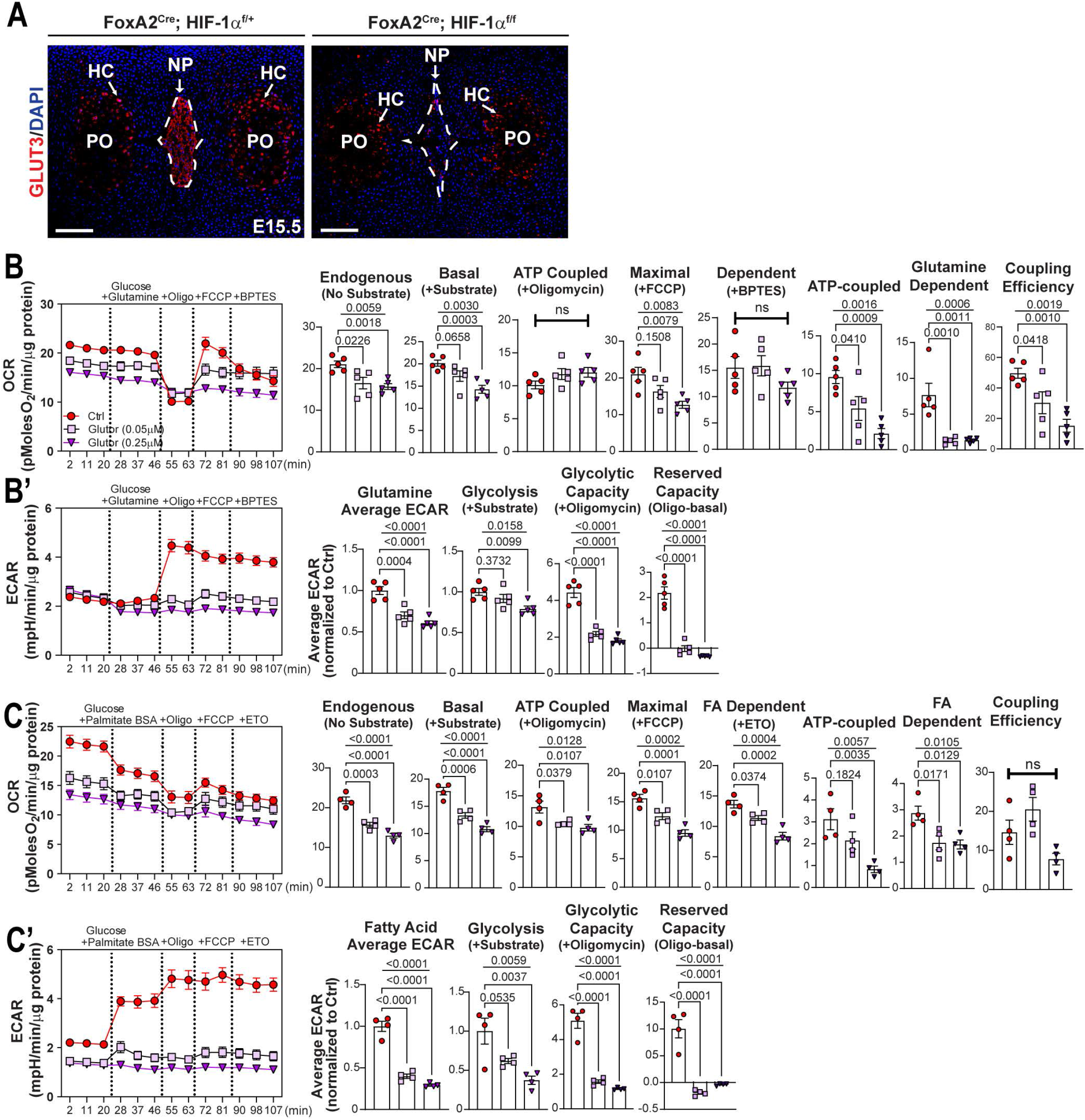
NP cells do not switch to glutamine utilization and fatty acid oxidation following loss of GLUT1 and GLUT3. (A) Representative image of GLUT3 staining in HIF-1α^f/f^ (WT) and HIF-1α^cKO^ (Foxa2^Cre^; HIF-1α^f/f^) mice (scale bar = 100 μm). PO: primary center of ossification, HC: hypertrophic chondrocytes. (B, B’) OCR and ECAR traces in Glutor treated and control cells in presence of Glutamine, and BPTES (B) quantification of endogenous, basal, maximal, and glutamine dependent OCR derived from traces shown in B, and (B’) glutamine-dependent average ECAR and glycolysis derived from traces in B’. (C, C’) OCR and ECAR traces in Glutor treated and control NP cells in presence of Palmitate-BSA, and ETO and (C) quantification of endogenous, basal, maximal, glutamine dependent OCR from traces in C, and (C’) average ECAR and glycolysis derived from traces shown in C’. Quantitative measurements represent mean ± SEM (n = 4-6 biological replicates, 4 technical replicates/experiment/group). Significance of differences was determined using one-way ANOVA with Sidak’s or Dunnett’s post hoc test as appropriate.

### NP cells do not switch to glutamine and fatty acid oxidation under glucose limiting conditions following GLUT1/3 inhibition

Since glucose is the major energy source for most cell types, it is plausible to consider that blocking its availability will shift cellular metabolism towards other substrates such as glutamine and/or fatty acids to maintain TCA cycle flux and ATP generation. Our previous studies have shown that although NP cells do not rely on oxidative phosphorylation for ATP production, the TCA cycle is intact and serves as a hub for generation of metabolic intermediates used in broad biosynthesis reactions. In recent years, TCA cycle intermediates have also been shown to “moonlight” in the nucleus where they engage in the epigenetic regulation of DNA and histone modifications (38).

To gain an insight into whether the significant loss of glucose import through GLUT1 and GLUT3 upregulates 1) glutamine metabolism or 2) fatty acid oxidation, we performed substrate dependent Seahorse experiments with glucose + glutamine or BSA-palmitate. To assess effects on glutamine metabolism, NP cells were treated with Glutor for 24 hours prior to Seahorse assessment. We analyzed raw OCR and ECAR traces beginning with endogenous conditions (no substrate), followed by the sequential addition of glucose + glutamine (basal), oligomycin (ATP synthase inhibitor), FCCP (oxidative phosphorylation uncoupler), and BPTES (glutaminase inhibitor) (Figure 4, B and B’). Glutor-treatment significantly decreased endogenous OCR and basal OCR in the presence of glucose and glutamine, implying glutamine oxidation alone is not sufficient to rescue OCR in NP cells (Figure 4B). Subsequently, control and Glutor-treated NP cells showed an expected decrease in OCR with oligomycin treatment, however FCCP was unable to shift 0.25 µM Glutor-treated cells to maximal OCR regardless of glutamine availability (Figure 4B). To determine the contribution of glutamine oxidation, we inhibited glutaminase with BPTES. Interestingly, BPTES treatment did not decrease OCR in the control group suggesting that NP cells do not prefer glutamine oxidation in presence of glucose. Importantly, taken together this data indicates that blocking glucose import through GLUT1/3 did not result in increased glutamine oxidation in NP cells (Figure 4B). Similarly, in the presence of glucose and glutamine, Glutor-treated cells showed a decrease in average ECAR, whereas, glycolysis-dependent ECAR was only affected at higher Glutor concentrations. Moreover, unlike control cells, Glutor-treated cells in presence of oligomycin (ETC inhibition) were unable to increase ECAR (Figure 4B’). Noteworthy, the presence of glutamine affected glucose-dependent ECAR in control cells i.e. glucose addition did not increase ECAR suggesting slowed glycolytic flux. These data imply that glutamine is not sufficient to rescue cell metabolism in the absence of glucose and in fact may interfere with glucose utilization by NP cells.

Considering that GLUT1/3 inhibition did not cause NP cells to switch to glutamine oxidation, we investigated whether glucose restriction resulted in the utilization of fatty acids to maintain TCA cycle flux. For these experiments, OCR was measured during endogenous conditions (no substrate), followed by the sequential addition of substrate (glucose + palmitate-BSA), oligomycin, FCCP, and finally Etomoxir (CPT1 inhibitor; inhibits mitochondrial import of fatty acid) (Figure 4, C and C’). Predictably, the endogenous OCR was decreased in Glutor-treated cells (Figure 4C) and the addition of glucose + palmitate-BSA did not rescue OCR (Figure 4C). Moreover, inhibiting fatty acid import with Etomoxir resulted in ∼25% decrease in OCR in control cells and ∼15-25% decrease in Glutor treated cells (Figure 4C). It was interesting to note that unlike glutamine, inclusion of BSA-palmitate with glucose did not interfere with ECAR induction in control cells. However, palmitate did not increase ECAR in Glutor treated cells, which remained significantly lower than control cells (Figure 4C’). Taken together, these metabolic experiments suggested that loss of GLUT1/3 function did not result in NP cells switching to glutamine or fatty acid oxidation and that glucose import through GLUT1/3 is indispensable for their metabolism.

### Nucleus pulposus-specific deletion of GLUT1 in skeletally mature mice does not affect disc health

To elucidate the role and test the apparent redundancy of GLUT1 seen in our *in vitro* experiments, we generated a conditional Glut1 knock-out in the mouse NP, driven by a tamoxifen-inducible K19^CreERT^ allele (Figure 5A). The K19^CreERT^;Glut1^f/f^ (cKO^K19^) and littermate control Glut1^f/f^ (WT) mice were injected with tamoxifen at 3 months (3M) and tissues collected at 9 months (9M). Robust *Glut1*/*Slc2a1* mRNA knockdown ∼80-90% was confirmed by qRT-PCR (Figure 5B). Deletion of GLUT1 protein was confirmed by fluorescence immunohistochemistry in both lumbar and caudal discs (Figure 5, C and C’) and well as Western blot (Figure 5, D and D’, Supplemental Figure 3, A-A’’). These results validate the cKO^K19^ mouse model and confirmed NP-specific deletion of GLUT1 protein expression in skeletally mature mice.

**Figure 5.**
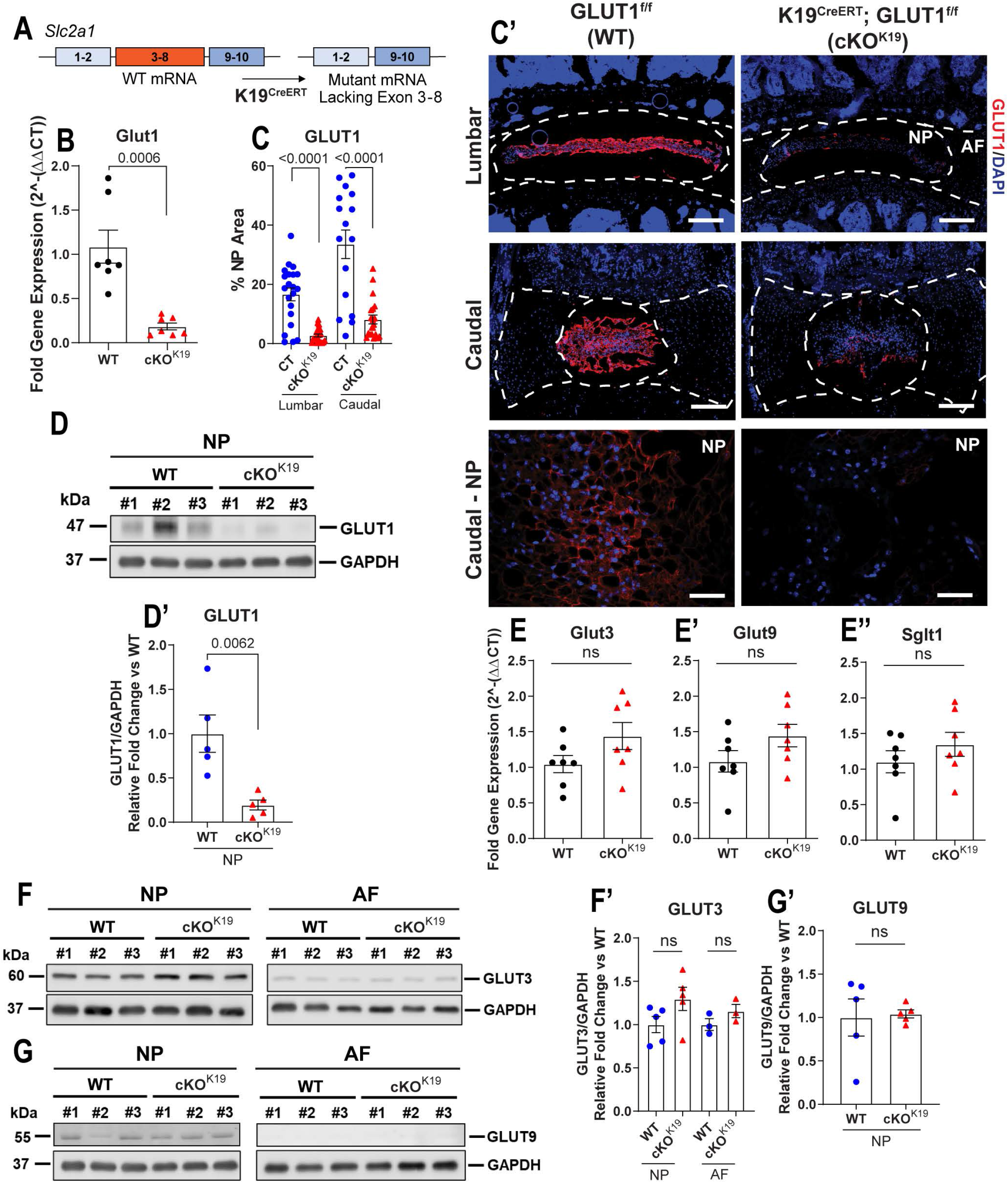
Conditional deletion of Glut1 in NP compartment of adult mice. (A) Schematic showing K19^Cre^ mediated deletion of *Glut1/Slc2a1* exon 3-8 to generate NP specific *Glut1* mutant. (B) quantification of *Glut1* in Glut1cKO^K19^ mice (n = 7 mice/genotype; 20 discs/animal). (C, C’) Representative IHC images and quantification of GLUT1 in 9-month-old WT and Glut1cKO^K19^ (scale bar = 200 μm and 50 μm) (n = 8 WT, 7 cKO mice; 6 lumbar and 3 caudal discs/animal). (D, D’) Western Blot and quantification of GLUT1 levels in NP of WT and Glut1cKO^K19^ mice (n = 5 mice/genotype; 20 discs/animal). White dotted lines demarcate disc compartments. (E-E’’) qRT-PCR of *Glut3, Glut9, Sglt1* in WT and Glut1cKO^K19^ mice (n = 7 mice/genotype; 20 disc/animal). (F, F’) Western blot and quantification of GLUT3 and (G, G’) GLUT9 (n = 5 mice/genotype; 20 disc/animal). Quantitative measurements represent mean ± SEM, significance of differences was determined using Mann-Whitney test.

We also confirmed the expression of alternative glucose transporters and determined if their levels are affected in Glut1 cKO^K19^ mice. There were comparable levels of *Glut3* and *Glut9*, as well as sodium-glucose cotransporter *Sglt1* mRNA between the WT and Glut1 cKO^K19^ mice (Figure 5, E-E’’). Furthermore, in line with previous findings in the human disc, Western blot analysis confirmed that GLUT3 and GLUT9 protein levels were robust in the NP of WT and cKO mice, with significantly lower abundance in the AF (Figure 5, F-G’, Supplemental Figure 3, B-C’’) (23). Based on these findings, we hypothesized that baseline levels of alternative glucose transporters in the NP, including GLUT3, GLUT9, and SGLT1 may be sufficient to sustain glucose transport and preserve disc health in GLUT1-deficient mice.

Histological changes were assessed in lumbar and caudal discs of WT and Glut1 cKO^K19^ at 9M of age (Figure 6A). Despite the robust expression of GLUT1 in the NP of healthy discs, Glut1 cKO^K19^ animals presented with no significant changes in the average grade of degeneration as measured by Modified Thompson Score (Figure 6B) or in the distribution in grades of degeneration (Figure 6B’) in the NP of lumbar discs. There was a change in the distribution in grades of degeneration in the lumbar AF, however it did not indicate increased degeneration. Curiously, caudal Glut1 cKO^K19^ discs showed a slight decrease in the average grade of NP degeneration (Figure 6B’’) and distributions skewed towards lower grades of degeneration for NP and AF as well (Figure 6B’’’). Taken together, NP-specific deletion of GLUT1 in adult mice had few discernible effects on disc degeneration in lumbar or caudal discs.

**Figure 6.**
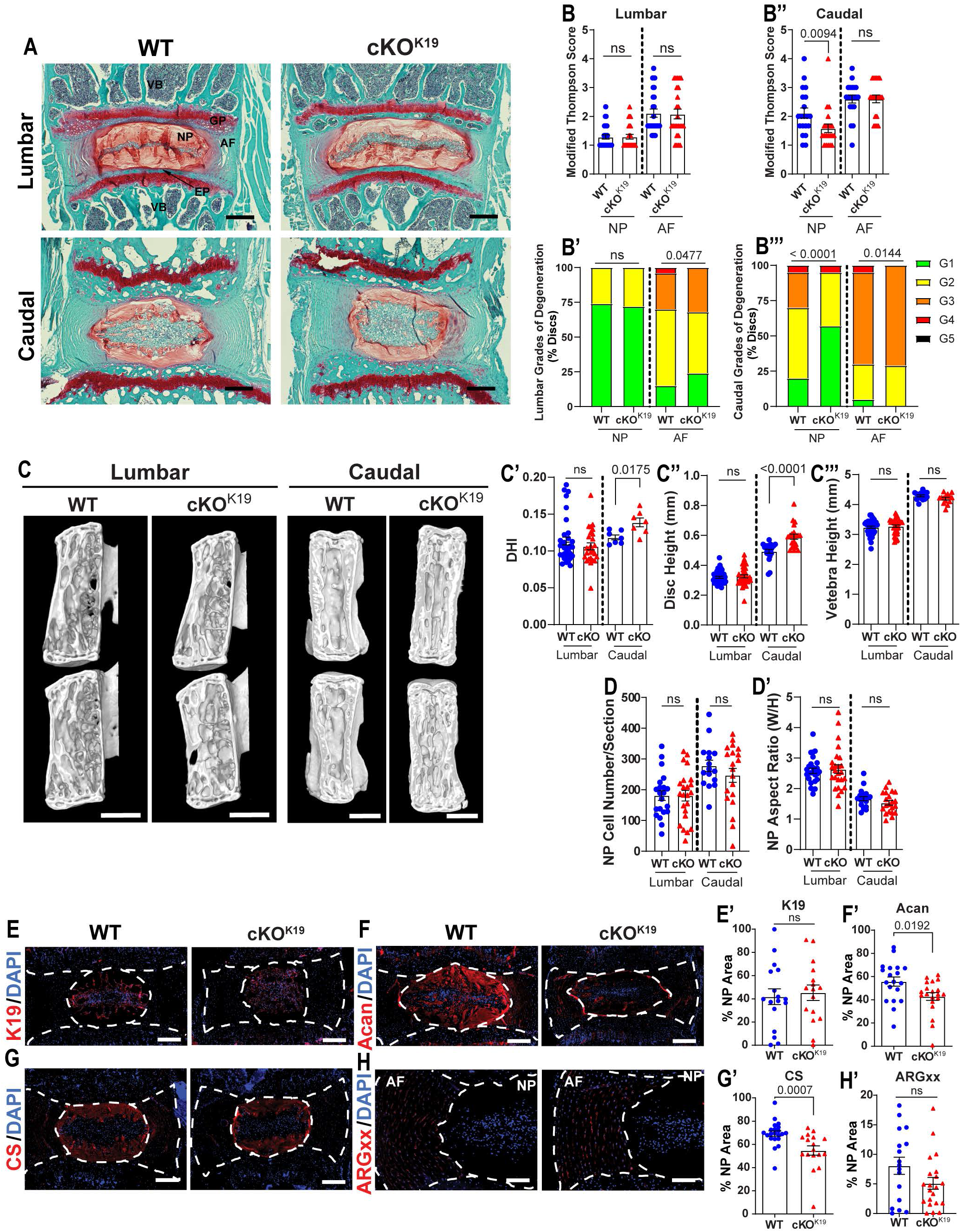
Glut1cKO^K19^ mice do not show adverse effects on intervertebral disc health. (A) Representative SafraninO/FastGreen images of 9-month-old WT and Glut1cKO^K19^ lumbar and caudal discs (scale bar = 200 μm). (B-B’’’) Modified Thompson Scores of NP and AF compartments in WT and Glut1cKO^K19^ lumbar and caudal discs. (n = 7 mice/genotype; 3-4 lumbar and 2-3 caudal discs/animal, 25-27 lumbar and 20-21 caudal discs/genotype) (C) Representative μCT images of lumbar and caudal motion segments (scale bar = 1 mm). (C’, C’’) Quantification of lumbar and caudal DHI, DH, and VBH. (n = 7 animals/genotype; Vertebrae L3-6, Ca5-6, and Discs L3/4-L6/S1, Ca5/6-7/8 per animal). (D, D’) Quantification of NP cell number and aspect ratio from lumbar and caudal discs. (n = 7 mice/genotype; 1-4 lumbar discs/animal, 2-3 caudal discs/animal; 25-27 lumbar and 20-21 caudal discs/genotype). (E-H’) Representative IHC and quantification of keratin-19 (K19), aggrecan (Acan), ARGxx, and chondroitin sulfate (CS) (scale bar = 200 μm, ARGxx scale bar = 100 μm). (n = 7 mice/genotype; 1-3 caudal discs/animal; 15-20 caudal discs/genotype). White dotted lines demarcate disc compartments. Significance for grading distribution was determined using a χ^2^ test. Significance of differences was determined using an unpaired t-test (C’, C’’’, D, D’, E’) or Mann-Whitney test (B, B’, C’-C”‘, D’, F’, G’, H’), as appropriate. Quantitative measurements represent mean ± SEM.

To evaluate the impact of the loss of GLUT1 on disc height (DH) and the disc height index (DHI), μCT imaging was performed on 9M WT and Glut1 cKO^K19^ mice (Figure 6C). In line with the histological analysis, we observed no changes in either DH or DHI in lumbar discs, however, there was a slight increase in both metrics in caudal discs (Figure 6C’-C’’’). The increased DHI in caudal discs could not be accounted for by changes in NP cell area (Figure 6D) or in the aspect ratio of the NP tissue compartment (Figure 6D’).

To determine whether the Glut1 cKO^K19^ show alterations in cell phenotypic makers and extracellular matrix (ECM) composition that do not reflect in histological grades of disc degeneration, we assessed the abundance of key extracellular matrix proteins in WT and Glut1 cKO^K19^ discs. Keratin-19 (K19), an NP cell phenotypic marker, showed no difference in abundance, suggesting the NP cell phenotype was maintained in Glut1 cKO^K19^ mice (Figure 6, E and E’). Interestingly, in the NP compartment of Glut1 cKO^K19^ mice, there was a significant decrease in the level of aggrecan – the major NP proteoglycan as well as chondroitin sulfate (CS) (Figure 5, F, F’, G, and G’). To understand if the decrease in aggrecan and its predominant glycosaminoglycan (GAG) chain was due to aggrecan turnover, we quantified changes in a neoepitope marker of cleaved aggrecan, ARGxx. Since there was no difference in ARGxx levels in discs of WT and Glut1 cKO^K19^ mice the results suggested a possible decrease in the synthesis of these molecules (Figure 5, H and H’). Accordingly, while adult Glut1 cKO^K19^ mice showed negligible differences in overall disc morphology, alteration in GAG synthesis suggests a small deficit in glucose availability and increased glucose flux into lactate/pyruvate to maintain ATP levels likely affecting hexosamine biosynthetic pathway.

We also made note of the significant difference in the distribution of grades of degeneration in the AF compartment of both lumbar and caudal discs in Glut1 cKO^K19^ mice (Figure 6, B’ and B’’’). To assess AF ECM, we analyzed collagen fiber architecture using Picrosirius Red staining coupled with polarized light imaging (Supplemental Figure 4, A and B). Under polarized light, green fluorescing fibers are thin, yellow fluorescing fibers are intermediate, and red fluorescing fibers are thick. We found that collagen fiber thickness is unaltered in lumbar AF (Supplemental Figure 4, A’ and A’’), however in Glut1 cKO^K19^ caudal discs, there was a significant increase in the percentage of thin (green) fibers and a decrease in thick (red) fibers (Supplemental Figure 4, B and B’’). This suggests that NP-specific deletion of GLUT1 may alter collagen fibers in the AF leading to a thinner and more immature fiber composition.

### Loss of GLUT1 does not alter expression of genes involved in compensatory metabolic pathways

To understand if disc health in Glut1 cKO^K19^ mice is maintained by regulation of compensatory mechanisms, we analyzed the NP transcriptome using microarray. There was a similar variance between the gene expression values in NP cells from WT and Glut1 cKO^K19^ mice, and the two phenotypes did not cluster independently along three principal components (Supplemental Figure 5A). When analyzed using a p-value cutoff of p ≤ 0.05 and a fold-change of ±2, several differentially expressed genes emerged from the dataset (Supplemental Figure 5, B and C), the most significant being *Slc2a1*, i.e. *Glut1*. However, PANTHER gene ontological analysis discovered no significantly enriched biological processes or molecular functions in the up- and down-regulated gene sets (data not shown). Furthermore, when the data were analyzed using more stringent parameters – FDR ≤ 0.05 and fold-change of ± 2 – the only differentially expressed gene was *Glut1*/*Slc2a1* (Supplemental Figure 5D).

The microarray analysis clearly shows that loss of GLUT1 causes strikingly few transcriptional changes in NP cells. This finding suggests that compensatory metabolic pathways do not require alterations in gene expression, despite the strict dependence of NP cells on glucose uptake. Changes in flux through alternative glucose transporters may therefore be sufficient to maintain the glycolytic capacity in the NP of GLUT1 cKO mice.

### GLUT1 expression in notochord/NP is not required for normal disc development

Considering that GLUT1 deletion in skeletally mature mice had no consequential effect on intervertebral disc health, we tested whether there was temporal dependency on GLUT1 function, specifically if it was required for normal embryonic and perinatal development of the NP. We crossed Glut1^f/f^ mice with notochord and floorplate specific constitutive Foxa2^Cre^ mice to generate notochord/NP specific GLUT1 knockout mice - Foxa2^Cre^; Glut1^f/f^ (i.e. cKO^Foxa2^) and littermate control (WT) mice (Figure 7A) were aged to postnatal day 7 (p7) and 14 weeks (14wk) where their morphology was assessed (Figure 7, B and C).

**Figure 7.**
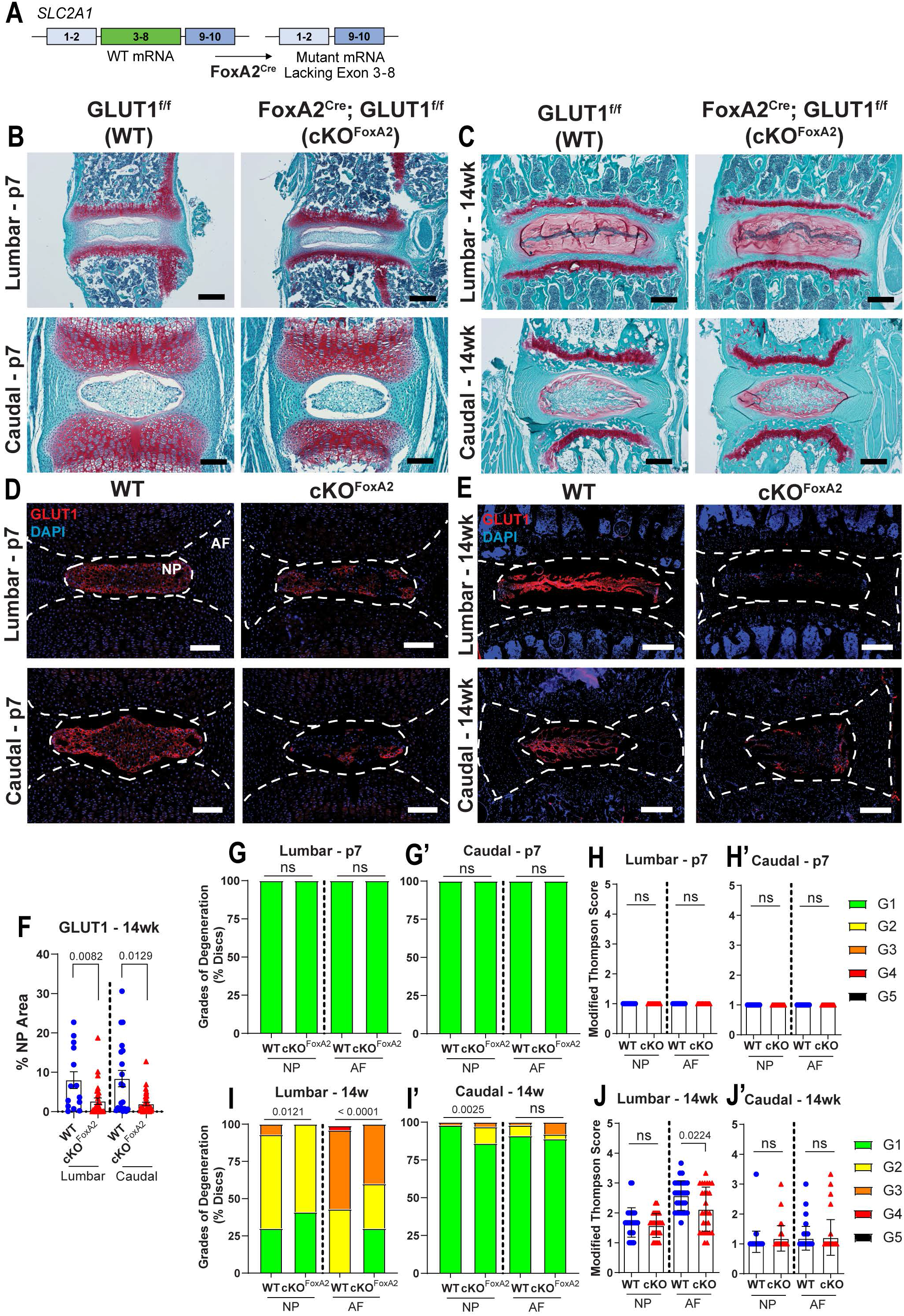
Glut1cKO^Foxa2^ mice do not show compromised disc health. (A) Schematic of Foxa2Cre mediated deletion of exons 3-8 in Glut1^f/f^ mice to generate a NP specific Glut1 cKO mice. (B-E) Representative images of (B, C) SafraninO/FastGreen staining and (D, E) GLUT1 IHC on Glut1^f/f^ (WT) and Foxa2^Cre^; Glut1^f/f^ (cKO^Foxa2^) discs at P7 and 14-weeks-old (P7 scale bar = 50 μm, 14-weeks scale bar = 200 μm). White dotted lines demarcate disc compartments. (F) Quantification of GLUT1 IHC at 14-weeks (n = 5 WT and 9 cKO^Foxa2^; 3 lumbar and 4 caudal discs/animal). (G, G’) Modified Thompson grade distribution and (H, H’) average grade scores of WT and cKO^Foxa2^ discs at P7 (n = 5 mice/genotype; 6 discs: 3 lumbar and 3 caudal/animal). (I, I’) Modified Thompson grade distribution and (J, J’) average grade scores of WT and cKO^Foxa2^ discs at 14-week (n = 10 WT and 10 cKO^Foxa2^; 3 lumbar and 4 caudal/animal). Significance for grade distribution was determined using a χ^2^ test. Significance of differences in Av. grade scores and % staining area was determined using Mann–Whitney test. Quantitative measurements represent mean ± SEM.

Immunohistochemistry confirmed a significant decrease in GLUT1 expression in the NP compartment of cKO^Foxa2^ mice at p7 (Figure 7D) and 14wk (Figure 7E). Despite the substantial GLUT1 knockdown (Figure 7F), lumbar and caudal discs from cKO^Foxa2^ did not present with any of the criteria for degeneration – in fact all NP and AF compartments from p7 mice had a Modified Thompson Score of 1 (Figure 7G-H’). At 14wk, the distribution in the grades of degeneration in cKO^Foxa2^ was different from the WT in lumbar NP and AF, as well as caudal NP, however the distributions favored lower grades of degeneration in the cKO^Foxa2^ mice (Figure 7, I and I’). Consequently, cKO^Foxa2^ mice lumbar AF had a slightly lower Modified Thompson Score, implying better morphological attributes, while lumbar NP and caudal NP and AF showed no significant difference from WT (Figure 7, J and J’). These results suggest that glucose uptake through GLUT1 is not a critical player for the development and early maintenance of the NP.

## DISCUSSION

The fact that loss of GLUT1 expression in the disc does not result in disc degeneration, transcriptomic changes, or metabolic disruption was baffling; especially considering the prominence of GLUT1 as a highly enriched NP phenotypic marker and the requirement for GLUT1 in the functional maintenance of other skeletal tissues including bone and cartilage (27, 28, 31). Indeed, our study underscores that although glucose is the indispensable metabolite, it appears that GLUT1 is not singularly required to maintain glycolytic capacity, and therefore, is not necessary for NP cell survival. Conversely, we hypothesize that maintaining ATP levels through glycolysis and TCA metabolites is so vital for disc health and NP cell viability that there is intrinsic redundancy to ensure an uninterrupted supply of glucose through glucose importers other than GLUT1.

In the hypoxic niche of the intervertebral disc, NP cells primarily rely on glycolysis for bioenergetics and highly express GLUT1 which is considered one of their key phenotypic markers. We therefore investigated the role GLUT1 plays in NP cell metabolism by using two highly potent GLUT1 inhibitors (35, 36). Surprisingly, since blocking GLUT1 in NP cells did not affect glucose uptake and the bioenergetic status of cells, the results suggested an alternate mechanism for glucose import. This finding was in contrast to bone cells that primarily depend on glucose for both differentiation and maturation during development and highly express GLUT1 which is responsible for 75% glucose uptake (27). One obvious explanation for this difference is that we and others have previously shown that NP cells express GLUT3 and GLUT9 (8, 23). Using a dual GLUT1/3 inhibitor Glutor, we determined that 80-90% glucose was imported through these two carriers (39). Importantly, unlike GLUT1 inhibition, Glutor treatment significantly inhibited glycolysis and diminished ATP production rate suggesting that GLUT3 is the critical high-capacity glucose importer in NP cells (Supplemental Figure 6). This also implies that NP cells in their avascular and diffusionally limited environment are intrinsically geared to import and utilize glucose as a metabolic substrate in a broad physiological concentration range given the diverse Km values for Glut1 (∼6.9 mM), Glut3 (1.8 mM) and Glut9 (0.61 mM) (40, 41).

It is known that under nutritional constraints or increased metabolic demand cellular function and survival is mediated by metabolic reprogramming (30, 42, 43). When glucose uptake is blocked, primarily glycolytic proinflammatory macrophages and cancer cells switch their metabolism to alternative substrates such as glutamine and fatty acids (30, 42). Similarly, a very recent study showed that deletion of GLUT1 in articular and growth plate chondrocytes resulted in increased cellular glutamine oxidation for survival (31). Similarly, in absence of glucose, osteoblasts shift to glutamine but not palmitate oxidation, whereas, myoblasts prefer palmitate oxidation over glutamine oxidation (27). It is important to note that NP cells also exhibit metabolic plasticity. For example, when lactate export through monocarboxylate transporter MCT4 is inhibited in NP cells, the cells undergo an incomplete metabolic switch from glycolytic to oxidative metabolism fueled by pyruvate oxidation (16). However, blocking lactate export still results in a 2-fold increase in the glycolytic intermediate, glucose-6-phosphate, implying glucose was still the major metabolite fueling pyruvate metabolism (16). While NP cells do have the capacity to metabolize alternative energy sources, such as fatty acids through mitochondrial β-oxidation (8), many studies including our own show that glucose starvation leads to cell death (12, 14, 22). Therefore, alternative energy sources such as glutamine, glycogen and fatty acids cannot compensate the loss of glucose import into NP cells. Furthermore, in support of this notion, we noted that when glucose import was impeded by GLUT1/3 inhibition, NP cells did not shift to glutamine or palmitate oxidation (Supplemental Figure 6). Rather, in contrast to other cell types, the overall oxygen consumption rate significantly decreased underscoring the fact that glucose is the primary metabolic fuel for ATP generation through glycolysis and is critical for NP cell survival (8, 39).

We have previously shown that in SM/J mice, a mouse model of early onset spontaneous degeneration, GLUT1 levels in the NP decline and as our current studies show, levels are lower during aging, an important risk factor associated with disc degeneration (33, 44, 45). These findings seemingly contradict our *in vitro* findings that showed minimal or no change in NP cell metabolism and their survival following GLUT1 inhibition. To clarify this apparent conflict and to delineate whether loss of GLUT1 expression *in vivo* causes disc degeneration, we utilized two mouse models, an inducible Glut1 cKO^K19^ and constitutive Glut1 cKO^Foxa2^ to determine a causal link between GLUT1 and intervertebral disc health. While there were some differences in grades of degeneration, between these mice the GLUT1 cKO^K19^ mice rather showed slightly lower grades of degeneration. However, that is not to say that GLUT1 loss-of-function is protective in the disc, rather the discs are generally healthy in both WT and cKO animals. In a parallel study, we used a constitutive GLUT1 cKO model to determine if GLUT1 is involved in disc development and post-natal maturation. This hypothesis stems from the idea that GLUT1 may play a key role in the HIF-1α-dependent glycolytic metabolism in the developing embryonic NP (9). The lack of apparent developmental defects in the discs from these mice further suggested that NP cells are refractory to loss of GLUT1 and that avenues of compensation are available to NP cells to allow them to survive the deletion of an integral glycolytic component. However, it may be of interest to ascertain whether functional compensation of GLUT1 or GLUT3 occurs when additional systemic stressors such as acute injury or aging are involved. Importantly, however, these observations raised an interesting prospect that a robust decrease in GLUT3 levels along with GLUT1 may contribute to metabolic failure and massive cell death of NP cells seen in HIF-1α^cKO^ mice by birth (9). Likewise decline in both GLUT3 and 1 may exacerbate metabolic restriction on cells in the aging disc in part driving loss of cells and an overall decreased biosynthesis of matrix molecules leading to age-dependent degeneration (1).

Another key observation was that despite GLUT1 loss-of-function, compensatory pathways do not involve transcriptional upregulation of other glucose transporters or metabolic enzymes. In fact, microarray analysis of NP from GLUT1 cKO^K19^ mice when analyzed with an adjusted FDR p-value of 0.05, failed to reveal a single differentially regulated transcript besides *Glut1/Slc2a1*. These data are corroborated by lack of changes in mRNA expression or protein levels of other *SLC2* family members and glucose importers expressed in the disc, namely GLUT3 and GLUT9 (23). Importantly, however, glucose import may be maintained in NP cells by increased flux through these SLC2 transporters without concomitant elevation in expression levels. Furthermore, glucose may also enter the NP cell through sodium-glucose cotransport (SGLT) of which six isoforms have been identified (46). SGLT is driven by the active extrusion of intracellular sodium, facilitating a concomitant import of extracellular glucose against plasma-membrane concentration gradients. Although SGLTs are attributed to glucose import across apical membranes, it is very possible they play a role in the NP compartment which is characterized by its high sodium concentrations in a hyperosmolar niche. While the contribution of these transporters has not yet been studied in the context of the intervertebral disc, it is likely to be secondary to SLC2 family members.

Taken together, our study provides the first evidence of functional redundancy in GLUT transporters in a physiologically hypoxic NP compartment of the intervertebral disc and highlights its uniquely different niche than other skeletal tissue like articular and growth plate cartilage.

Importantly, our findings underscore the importance of glucose as the indispensable metabolic fuel for NP cells and provide a vital baseline for any cell-based therapies aimed at restoring the function of the degenerating disc.

## METHODS

### Mice

All procedures regarding collection of animal tissues were performed as per approved protocols by the Institutional Animal Care and Use Committee (IACUC) of Thomas Jefferson University, in accordance with the IACUC’s relevant guidelines and regulations. For postnatal deletion in NP compartment, *Glut1*^*f/f*^ mice (47) were crossed with K19^CreERT^ mice (*Krt19*^*tm1(cre/ERT)Ggu*^/J, Jackson Stock # 026925) to produce K19^CreERT^; Glut1^f/f^ (cKO^K19^) mice, Glut1^f/f^ littermate mice served as control (WT), and injected with three consecutive Tamoxifen at 3-months at a 100 μg/g.b.w.; these mice were collected 6-months post-recombination (9-months-old) (n = 8 *Glut1*^*f/f*^ mice: 5 males, 3 females and 7 cKO^K19^ mice: 3 males, 4 females) (48). For deletion of GLUT1 at embryonic time points, Glut1^f/f^ mice were crossed with Foxa2^Cre^ mice to produce Foxa2^Cre^; Glut1^f/f^ (cKO^FoxA2^) mice, littermate Foxa2^Cre^; Glut1^f/+^ and Glut1^f/f^ mice served as control (WT); these mice were assessed at P7 (n = 5 mice/genotype), 14-weeks (n = 10 mice/genotype; 5 female WT, 5 male WT, 4 female cKO^FoxA2^, 6 male cKO^FoxA2^). Foxa2^Cre^ allele drives robust expression specifically in the notochord and floorplate using combination of 5’ notochord and 3’ floorplate enhancers under the control of *Hspa1* promoter (49).

### Immunofluorescence Microscopy and digital image analysis

7-μm thick mid-coronal disc sections were de-paraffinized and incubated in microwaved citrate buffer for 20 min or proteinase K for 10 min at room temperature, or Chondroitinase ABC for 30 min at 37 °C for antigen retrieval. Appropriate wildtype and cKO histological sections were blocked in 5% normal serum (Thermo Fisher Scientific, 10000 C) in PBS-T (0.4% Triton X-100 in PBS) and incubated with antibody against KRT19 (1:3, DSHB, TROMA-III/supernatant), CA3 (1:150, Santa Cruz, sc-50715), Aggrecan (1:50, Millipore, AB1031), GLUT-1 (1:200, Abcam, ab40084), ARGxx (1:200, Abcam, ab3773) and GLUT3 (1:200, Proteintech, 20403-1-AP) in blocking buffer at 4°C overnight. For GLUT-1 (1:200, Abcam, ab40084) and ARGxx (1:200, Abcam, ab3773) staining, Mouse on Mouse Kit (Vector laboratories, BMK2202) was used for blocking and primary antibody incubation. Tissue sections were thoroughly washed and incubated with Alexa Fluor®594 (Ex: 591 nm, Em: 614 nm) conjugated secondary antibody (Jackson ImmunoResearch Lab, Inc.), at a dilution of 1:700 for 1 h at room temperature in dark. The sections were washed again with PBS-T (0.4% Triton X-100 in PBS) and mounted with ProLong® Gold Antifade Mountant with DAPI (Thermo Fisher Scientific, P36934). All mounted slides were visualized with Axio Imager 2 (Carl Zeiss) using 5×/0.15 NAchroplan (Carl Zeiss), 10×/0.3 EC Plan-Neofluar (Carl Zeiss), or 20x/0.5 EC Plan-Neofluar (Carl Zeiss) objectives, X-Cite® 120Q Excitation Light Source (Excelitas), AxioCam MRm camera (Carl Zeiss), or LSM800 (Carl Zeiss) 20x/0.8 or 40x/1.3 Oil Plan-Apochromat (Carl Zeiss), AxioCam 506 mono (Carl Zeiss), and Zen2TM software (Carl Zeiss). DAPI-positive cells were analyzed to assess cell number in disc compartments. All quantifications were done in 8-bit greyscale using the Fiji package of ImageJ. Images were thresholded to create binary images, and NP and AF compartments were manually segmented using the Freehand Tool. These defined regions of interest were analyzed either using the Analyze Particles (cell number quantification) function or the Area Fraction measurement.

### Protein extraction and Western Blotting

Mouse NP tissue was carefully separated from AF tissue under a dissecting microscope as described previously (50, 51, 52, 53, 54). Following NP tissue extraction from Wildtype and GLUT1 cKO^K19^ mice, cells were washed on ice with ice-cold 1X PBS with protease inhibitor cocktail (Thermo Scientific). Cell were lysed with lysis buffer containing 1X protease inhibitor cocktail (Thermo Scientific), NaF (4 mM), Na3VO4 (20 mM), NaCl (150 mM), β-glycerophosphate (50 mM), and DTT (0.2 mM). Total protein (35 µg per sample) was resolved on 10% SDS-polyacrylamide gels and transferred to PVDF membranes (Fisher Scientific).

Membranes were blocked with 5% nonfat dry milk in TBST (50 mM Tris pH 7.6, 150 mM NaCl, 0.1% Tween 20) and incubated overnight at 4°C in 5% nonfat dry milk in TBST with anti-GLUT3 (1:500, Proteintech, 20403-1-AP) or anti-GLUT9 (1:500, Abcam, ab223470) antibodies.

Specificity of all antibodies has been validated by the manufacturers using siRNA or negative control IgG. Immunolabeling was detected using ECL reagent and imaged using LAS4000 system (GE Life Sciences). Densitometric analysis was performed using ImageJ. All quantitative data is represented as mean ± SEM, n = 2-5 animal/genotype.

### Isolation of Rat NP cells, hypoxic culture, cell treatments

Rat NP cells were isolated as reported previously by Risbud and colleagues (5). Rat NP cells were maintained in Dulbecco’s Modification of Eagle’s Medium (DMEM) supplemented with 10% FBS and antibiotics. Cells were cultured in a Hypoxia Work Station (Invivo2 400, Ruskinn, UK) with a mixture of 1% O2, 5% CO2 and 94% N2. To investigate the effect of GLUT inhibition, NP cells were treated with 1) a cocktail of GLUT1 inhibitors (BAY-876, 0.01, 0.1 µM, Cayman, 19961) or (WZB-117, 1, 10 μM, Cayman, 19900) for 1 to 72 hours or 2) a GLUT1, 2 and 3 inhibitor (Glutor, 0.05, 0.1, and 0.25 µM, Sigma, SML2765) for 6 to 72 hours. Viability measurements following treatment of NP cells in hypoxia with BAY-876, WZB-117 and Glutor for 24-72 h were performed using a Calcein AM cell viability assay as per manufacturer’s instructions (Invitrogen, C3100MP). All *in vitro* experiments were performed in hypoxia (1% O2) at least 3-6 independent times with 4 replicates/experiment/group and data represented as mean ± SEM.

### Seahorse XF Analyzer Respiratory Assay

The oxygen ECAR and OCR were measured using method reported by Mookerjee and colleagues (37), briefly, rat NP cells were plated in 24-well Seahorse V7-PS assay plate and cultured for 36 hours in normoxia conditions. 24 h prior to assay cells were treated with GLUT1 inhibitors BAY-876 or WZB-117 and cultured under hypoxia. Prior to measurement cells were washed three times with 500 ul of KRPH (Krebs Ringer Phosphate HEPES) and incubated 37°C for 1 hour under 100% air. OCR and ECAR was measured by addition of 10 mM glucose, 2 μg oligomycin, 1 μM rotenone plus 1 μM myxothiazol. The rate of oxygen consumption and extracellular acidification were normalized to protein content of the appropriate well. For the substrate dependency assay, 24 h prior to assay, cells were treated with GLUT1/3 inhibitor Glutor and cultured under hypoxia. Prior to measurement cells were washed three times with 500 μl of KRPH (Krebs Ringer Phosphate HEPES) and incubated 37°C for 1 hour under 100% air. OCR and ECAR was measured by addition of 10 mM glucose plus 4mM glutamine, 2 μg oligomycin, 1 μM FCCP and 5 μM BPTES for glutamine oxidation. For fatty acid oxidation KRPH was supplemented with 0.5 mM L-carnitine and incubated 37°C for 1 hour under 100% air. OCR and ECAR was measured by addition of 10 mM glucose plus Palmitate-BSA (palmitate concentration 150 μM), 2 μg oligomycin, 1 μM FCCP and 5 μM Etomoxir. The rate of oxygen consumption and extracellular acidification were normalized to protein content of the appropriate well.

### 2-DG uptake assay

Rat NP cells (20,000 cells/well) were seeded in a 96 well plate in a complete medium and were grown for 48h in normoxia. After 48h cells were treated with glucose transporter inhibitors for 6 and 24 hours under hypoxia. The 2-DG uptake was measured following the (Abcam, ab136956) protocol. Briefly, adherent cells were washed three times with freshly prepared Krebs-Ringer-Phosphate-HEPES (KRPH) buffer (20mM HEPES, 5mM KH2PO4, 1mM MgSO4, 1mM CaCl2, 136mM NaCl, 4.7mM KCl, pH 7.4, 2% BSA). 2-DG (1μM) was added to the cells together with the respective inhibitors in KRPH buffer and incubated for 40 mins. DMSO and without 2-DG served as control. After 40 mins the 2-DG uptake was stopped by removing assay buffer and washing 3 times with ice-cold KRPH buffer. Cell lysis and endogenous NAD(P) degradation were performed by adding extraction buffer to the cells and incubated for 30 min at 85°C. Cell lysates were kept on ice for 5 min. and neutralization was performed by adding neutralization buffer. Then 2-DG uptake was measured following the protocol. All quantitative data are represented as mean ± SEM, n = 6 independent experiments, 4 replicates/experiment/group.

### Real Time RT-PCR Analysis

NP tissue was micro-dissected from 9M Wildtype and Glut1 cKO^K19^ animals. Tissues from L1/2-L6/S1 and Ca1/2-Ca14/15 of same mouse were pooled and served as a single sample stored into RNAlater® Reagent (Invitrogen, Carlsbad, CA) for minimum 2 days at -80°C (n = 7 mice/genotype; 20 discs/animal). Samples were homogenized with a Pellet Pestle Motor (Sigma Aldrich, Z359971), and RNA was extracted using RNeasy® Mini kit (Qiagen). RNA was quantified on a Nanodrop ND-100 spectrophotometer (Thermo Fisher Scientific). Purified RNA was then converted to cDNA using EcoDry™ Premix (Clonetech). Gene specific primers (IDT, IN) and the template cDNA were added to Power SYBR Green master mix (Applied Biosystems). Primer sets were designed and by IDT, Inc. (S. Table 1). Quantification of expression was done by the StepOnePlus Realtime PCR system (Applied Biosystem) using ΔΔCT method and *Hprt* to normalize gene expression.

### Mouse Histological analysis

Mouse spines were harvested and fixed in 4% PFA for 24-48 hours and decalcified in EDTA (12.5-20%) at 4°C for 15-21 days prior to paraffin embedding. Mid-coronal 7 μm disc sections (Ca5/6-Ca8/9, L1/2-L6/S1) were stained with Safranin O/Fast Green/Hematoxylin or Picrosirius red, then visualized using a light microscope (AxioImager 2, Carl Zeiss) or a polarizing microscope (Eclipse LV100 POL, Nikon). Histopathological grading was performed on n = 5 mice/genotype with 6 discs per mouse (30 discs/genotype) at P7 and 14 weeks (WT and Glut1cKO^Foxa2^); n = 8 mice/genotype with 6 discs per mouse (48 discs/genotype) at 9 months (WT and Glut1cKO^K19^). Modified Thompson grading was used to score NP and AF compartments by 3 blinded graders. Aspect ratio of NP was determined by width divided by height of the NP tissue measured from Safranin O/Fast Green staining images of mid-coronal tissue sections from 9 month-old WT and Glut1cKO^K19^ animals, n = 6 mice/genotype with 3 discs per mouse (18 discs/genotype) using ImageJ software (http://rsb.info.nih.gov/ij/).

### Micro-CT analysis

Micro-CT (μCT) scans (Bruker, Skyscan 1275) were performed on WT and Glut1cKO^K19^ spines fixed with 4% PFA. Lumbar spine segments incorporating L2-S1 (7 mice/genotype) were scanned with an energy of 50 kV at 200 μA, resulting in 15 μm^3^ voxel size resolution.

Intervertebral disc height and the vertebral length were measured and averaged along the dorsal, midline, and ventral regions in the sagittal plane and Disc height index (DHI) was calculated as previously described (44, 50, 51, 52, 55).

### Transcriptomic Analysis and Enriched Pathways

RNA was quantified on a Nanodrop ND-100 spectrophotometer, followed by RNA quality assessment analysis on an Agilent 2200 TapeStation (Agilent Technologies, Palo Alto, CA). Fragmented biotin labeled cDNA (from 100 ng of RNA) was synthesized using the GeneChip WT Plus kit according to ABI protocol (Thermo Fisher Scientific). Gene chips, Mouse Clariom S were hybridized with 2.5 μg fragmented and biotin-labeled cDNA in 100 μl of hybridization cocktail. Arrays were washed and stained with GeneChip hybridization wash & stain kit using Gene chip Fluidic Station 450. Chips were scanned on an Affymetrix Gene Chip Scanner 3000 7G, using Command Console Software. Quality Control of the experiment was performed by Expression Console Software v 1.4.1. Chp files were generated by sst-rma normalization from Affymetrix .cel file using Expression Console Software. Experimental group was compared with control group by using Transcriptome array console 4.0 and DEGs with a Fold Change ±2, p-value or FDR<0.05. PANTHER tool was used to compute enriched pathways in DEGs that are altered with a Fold Change ±2, p-value <0.05. For evaluating highly expressed genes in the developing notochord/NP and NP of healthy adult mice, data deposited by Peck *et al*.

GSE100934 (32) and Novais *et al*. GSE134955 (33) were used.

### Statistical analysis

Quantitative data are presented as mean ± SEM and data distribution was checked with Shapiro-Wilk normality test, and unpaired t-test or Mann-Whitney test was used as appropriate. Comparisons between more than 2 groups were performed by the one-way ANOVA or Kruskal-Wallis test with appropriate post-hoc analyses (Dunn’s or Dunnett’s or Sidak’s multiple comparisons test) using Prism9 (Graphpad Software); p<0.05. For histopathological analysis showing percent-degenerated-discs and Picrosirius red percentage AF area, χ2 test was used.

### Study Approval

All procedures regarding collection of animal tissues were performed as per approved protocols by the Institutional Animal Care and Use Committee (IACUC) of Thomas Jefferson University, in accordance with the IACUC’s relevant guidelines and regulations.

## Data Availability

All data generated or analyzed during this study are included in this published article (and its Supplementary files) or deposited on the GEO database (GSE208396).

## Acknowledgements

This work is supported by grants from the National Institutes of Health R01 AR055655, R01 AG073349, R01 AR074813 (MVR) and T32 AR052273 (IMS). We acknowledge Drs. E. Dale Abel for proving GLUT1^f/f^ mice, Ernestina Schipani for FoxA2^Cre^; HIF-1α^f/f^ mice and Michael Kuehn for FoxA2^Cre^.

## Authors Contributions

ESS, SNJ, MVR, IMS designed the project. SNJ, VM, ESS, DHN performed all experiments. ESS, SNJ, VM, IMS and MVR wrote and edited the manuscript.

## Disclosures

The authors declare that they have no conflicts of interest or disclosures with the contents of this article.

## Notes

### Competing Interest Statement

The authors have declared no competing interest.

### Summary of Updates

Added additional references and updated the methods section. Revised all units to be in uM. New data was added as Fig 2F-G'', Fig 3C-E, and Supplemental Figures 1D, 2, and 6. Overall, the number of the main figures increased from 6 to 7, and the supplemental figures increased from 4 figures and 1 table to 6 figures and 1 table.

## REFERENCES

1. Silagi ES, Schipani E, Shapiro IM, Risbud M V. The role of HIF proteins in maintaining the metabolic health of the intervertebral disc [Internet]. Nat. Rev. Rheumatol. 2021;17(7):426–439.

2. Tessier S, Madhu V, Johnson ZI, Shapiro IM, Risbud M V. NFAT5/TonEBP controls early acquisition of notochord phenotypic markers, collagen composition, and sonic hedgehog signaling during mouse intervertebral disc embryogenesis [Internet]. Dev. Biol. 2019;455(2):369–381.

3. Risbud M V. et al. Defining the phenotype of young healthy nucleus pulposus cells: Recommendations of the spine research interest group at the 2014 annual ORS meeting. J. Orthop. Res. 2015;33(3):283–293.

4. Rajpurohit R, Risbud M, Ducheyne P, Vresilovic E, Shapiro I. Phenotypic characteristics of the nucleus pulposus: expression of hypoxia inducing factor-1, glucose transporter-1 and MMP-2 [Internet]. Cell Tissue Res. 2002;308(3):401–407.

5. Risbud M V. et al. Nucleus pulposus cells express HIF-1 alpha under normoxic culture conditions: a metabolic adaptation to the intervertebral disc microenvironment.. J. Cell. Biochem. 2006;98(1):152–159.

6. Fujita N, Chiba K, Shapiro IM, Risbud M V. HIF-1alpha and HIF-2alpha degradation is differentially regulated in nucleus pulposus cells of the intervertebral disc. J Bone Min. Res 2012;27(2):401–412.

7. Madhu V et al. Hypoxic Regulation of Mitochondrial Metabolism and Mitophagy in Nucleus Pulposus Cells Is Dependent on HIF-1α–BNIP3 Axis. J. Bone Miner. Res. [published online ahead of print: 2020]; doi:10.1002/jbmr.4019

8. Agrawal A et al. Normoxic stabilization of HIF-1alpha drives glycolytic metabolism and regulates aggrecan gene expression in nucleus pulposus cells of the rat intervertebral disk.. Am. J. Physiol. Cell Physiol. 2007;293(2):C621–C631.

9. Merceron C et al. Loss of HIF-1alpha in the notochord results in cell death and complete disappearance of the nucleus pulposus. PLoS One 2014;9(10):e110768.

10. Schoepflin ZR, Silagi ES, Shapiro IM, Risbud MV. PHD3 is a transcriptional coactivator of HIF-1a in nucleus pulposus cells independent of the PKM2-JMJD5 axis. FASEB J. 2017;31(9). doi:10.1096/fj.201601291R

11. Grunhagen T, Shirazi-Adl A, Fairbank JCT, Urban JPG. Intervertebral Disk Nutrition: A Review of Factors Influencing Concentrations of Nutrients and Metabolites. Orthop. Clin. North Am. 2011;42(4):465–477.

12. Bibby SRS, Urban JPG. Effect of nutrient deprivation on the viability of intervertebral disc cells [Internet]. Eur. Spine J. 2004;13(8):695–701.

13. Naqvi SM, Buckley CT. Extracellular matrix production by nucleus pulposus and bone marrow stem cells in response to altered oxygen and glucose microenvironments. J. Anat. [published online ahead of print: 2015]; doi:10.1111/joa.12305

14. Guehring T et al. Notochordal intervertebral disc cells: sensitivity to nutrient deprivation. [Internet]. Arthritis Rheum. 2009;60(4):1026–34.

15. Hartman R et al. Age-dependent changes in intervertebral disc cell mitochondria and bioenergetics. Eur. Cells Mater. 2018;36:171–183.

16. Silagi ES et al. Lactate Efflux From Intervertebral Disc Cells Is Required for Maintenance of Spine Health. J. Bone Miner. Res. 2020;35(3):550–570.

17. Wang D et al. Lactate oxidative phosphorylation by annulus fibrosus cells: evidence for lactate-dependent metabolic symbiosis in intervertebral discs [Internet]. Arthritis Res. Ther. 2021;23(1). doi:10.1186/S13075-021-02501-2

18. Ashinsky BG et al. Intervertebral Disc Degeneration Is Associated With Aberrant Endplate Remodeling and Reduced Small Molecule Transport [Internet]. J. Bone Miner. Res. 2020;35(8):1572–1581.

19. Silagi ES et al. Bicarbonate Recycling by HIF-1–Dependent Carbonic Anhydrase Isoforms 9 and 12 Is Critical in Maintaining Intracellular pH and Viability of Nucleus Pulposus Cells. J. Bone Miner. Res. 2018;33(2):338–355.

20. Wang C, Gonzales S, Levene H, Gu W, Huang CYC. Energy metabolism of intervertebral disc under mechanical loading. J. Orthop. Res. 2013;31:1733–8.

21. Bartels EM, Fairbank JC, Winlove CP, Urban JP. Oxygen and lactate concentrations measured in vivo in the intervertebral discs of patients with scoliosis and back pain.. Spine (Phila. Pa. 1976). 1998;23(1):1–7; discussion 8.

22. Bibby SRS, Jones D a, Ripley RM, Urban JPG. Metabolism of the intervertebral disc: effects of low levels of oxygen, glucose, and pH on rates of energy metabolism of bovine nucleus pulposus cells. [Internet]. Spine (Phila. Pa. 1976). 2005;30(5):487–496.

23. Richardson SM, Knowles R, Tyler J, Mobasheri A, Hoyland JA. Expression of glucose transporters GLUT-1, GLUT-3, GLUT-9 and HIF-1alpha in normal and degenerate human intervertebral disc. [Internet]. Histochem. Cell Biol. 2008;129(4):503–11.

24. Heilig C et al. Implications of glucose transporter protein type 1 (GLUT1)-haplodeficiency in embryonic stem cells for their survival in response to hypoxic stress [Internet]. Am. J. Pathol. 2003;163(5):1873–1885.

25. Wang D et al. A mouse model for Glut-1 haploinsufficiency. Hum. Mol. Genet. [published online ahead of print: 2006]; doi:10.1093/hmg/ddl032

26. Swarup A et al. Modulating GLUT1 expression in retinal pigment epithelium decreases glucose levels in the retina: impact on photoreceptors and Müller glial cells [Internet]. Am. J. Physiol. Cell Physiol. 2019;316(1):C121–C133.

27. Wei J et al. Glucose Uptake and Runx2 Synergize to Orchestrate Osteoblast Differentiation and Bone Formation [Internet]. Cell 2015;161(7):1576–1591.

28. Lee SY, Abel ED, Long F. Glucose metabolism induced by Bmp signaling is essential for murine skeletal development. Nat. Commun. 2018;9:4831.

29. Swarup A et al. Deletion of GLUT1 in mouse lens epithelium leads to cataract formation [Internet]. Exp. Eye Res. 2018;172:45–53.

30. Freemerman AJ et al. Myeloid Slc2a1-Deficient Murine Model Revealed Macrophage Activation and Metabolic Phenotype Are Fueled by GLUT1 [Internet]. J. Immunol. 2019;202(4):1265–1286.

31. Wang C et al. Deletion of Glut1 in early postnatal cartilage reprograms chondrocytes toward enhanced glutamine oxidation [Internet]. Bone Res. 2021;9(1). doi:10.1038/S41413-021-00153-1

32. Peck SH et al. Whole Transcriptome Analysis of Notochord-Derived Cells during Embryonic Formation of the Nucleus Pulposus [Internet]. Sci. Rep. 2017;7(1). doi:10.1038/S41598-017-10692-5

33. Novais EJ et al. Comparison of inbred mouse strains shows diverse phenotypic outcomes of intervertebral disc aging. Aging Cell 2020;19(5):213148.

34. Silagi ES, Batista P, Shapiro IM, Risbud MV. Expression of Carbonic Anhydrase III, a Nucleus Pulposus Phenotypic Marker, is Hypoxia-responsive and Confers Protection from Oxidative Stress-induced Cell Death. Sci. Rep. 2018;8(1):4856.

35. Wu Q et al. GLUT1 inhibition blocks growth of RB1-positive triple negative breast cancer [Internet]. Nat. Commun. 2020;11(1). doi:10.1038/S41467-020-18020-8

36. Liu Y et al. A small-molecule inhibitor of glucose transporter 1 downregulates glycolysis, induces cell-cycle arrest, and inhibits cancer cell growth in vitro and in vivo [Internet]. Mol. Cancer Ther. 2012;11(8):1672–1682.

37. Mookerjee SA, Gerencser AA, Nicholls DG, Brand MD. Quantifying intracellular rates of glycolytic and oxidative ATP production and consumption using extracellular flux measurements [Internet]. J. Biol. Chem. 2017;292(17):7189–7207.

38. Martínez-Reyes I, Chandel NS. Mitochondrial TCA cycle metabolites control physiology and disease [Internet]. Nat. Commun. 2020;11(1). doi:10.1038/S41467-019-13668-3

39. Reckzeh ES et al. Inhibition of Glucose Transporters and Glutaminase Synergistically Impairs Tumor Cell Growth [Internet]. Cell Chem. Biol. 2019;26(9):1214-1228.e25.

40. Burant CF, Bell GI. Mammalian facilitative glucose transporters: evidence for similar substrate recognition sites in functionally monomeric proteins [Internet]. Biochemistry 1992;31(42):10414–10420.

41. Doblado M, Moley KH. Facilitative glucose transporter 9, a unique hexose and urate transporter [Internet]. Am. J. Physiol. Endocrinol. Metab. 2009;297(4). doi:10.1152/AJPENDO.00296.2009

42. Brooks Robey R et al. Metabolic reprogramming and dysregulated metabolism: cause, consequence and/or enabler of environmental carcinogenesis? [Internet]. Carcinogenesis 2015;36 Suppl 1(Suppl 1):S203–S231.

43. Ito K, Suda T. Metabolic requirements for the maintenance of self-renewing stem cells [Internet]. Nat. Rev. Mol. Cell Biol. 2014;15(4):243–256.

44. Choi H et al. A novel mouse model of intervertebral disc degeneration shows altered cell fate and matrix homeostasis. Matrix Biol. 2018;70:102–122.

45. Ohnishi T et al. Caspase-3 knockout inhibits intervertebral disc degeneration related to injury but accelerates degeneration related to aging [Internet]. Sci. Rep. 2019;9(1). doi:10.1038/S41598-019-55709-3

46. Allison S. Structures of sodium-glucose cotransporters [Internet]. Nat. Rev. Nephrol. 2022;18(3):137.

47. Winkler EA et al. GLUT1 reductions exacerbate Alzheimer’s disease vasculo-neuronal dysfunction and degeneration [Internet]. Nat. Neurosci. 2015;18(4):521–530.

48. Mohanty S, Pinelli R, Dahia CL. Characterization of Krt19 CreERT allele for targeting the nucleus pulposus cells in the postnatal mouse intervertebral disc [Internet]. J. Cell. Physiol. 2020;235(1):128–140.

49. Uetzmann L, Burtscher I, Lickert H. A mouse line expressing Foxa2-driven Cre recombinase in node, notochord, floorplate, and endoderm [Internet]. Genesis 2008;46(10):515–522.

50. Tessier S et al. TonEBP-deficiency accelerates intervertebral disc degeneration underscored by matrix remodeling, cytoskeletal rearrangements, and changes in proinflammatory gene expression. Matrix Biol. 2020;87:94–111. doi: 10.1016/j.matbio.2019.10.007

51. Tsingas M et al. Sox9 deletion causes severe intervertebral disc degeneration characterized by apoptosis, matrix remodeling, and compartment-specific transcriptomic changes. Matrix Biol. 2020;94:110–133. doi: 10.1016/j.matbio.2020.09.003

52. Novais EJ et al. Long-term treatment with senolytic drugs Dasatinib and Quercetin ameliorates age-dependent intervertebral disc degeneration in mice. Nat Commun. 2021;12(1):5213. doi: 10.1038/s41467-021-25453-2

53. Piprode V et al. An optimized step-by-step protocol for isolation of nucleus pulposus, annulus fibrosus, and end plate cells from the mouse intervertebral discs and subsequent preparation of high-quality intact total RNA. JOR Spine 2020;3(3):e1108. doi: 10.1002/jsp2.1108

54. Kushioka J et al. A novel and efficient method for culturing mouse nucleus pulposus cells. Spine J. 2019;19(9):1573–1583. doi: 10.1016/j.spinee.2019.04.005

55. Tajerian M et al. DNA methylation of SPARC and chronic low back pain. Mol Pain. 2011;7:65. doi: 10.1186/1744-8069-7-65

