## Supplemental Figures and Table for "GLUT1 is redundant in hypoxic and glycolytic nucleus pulposus cells of the intervertebral disc"

**This PDF file includes:**

Figures S1 to S6
Tables S1

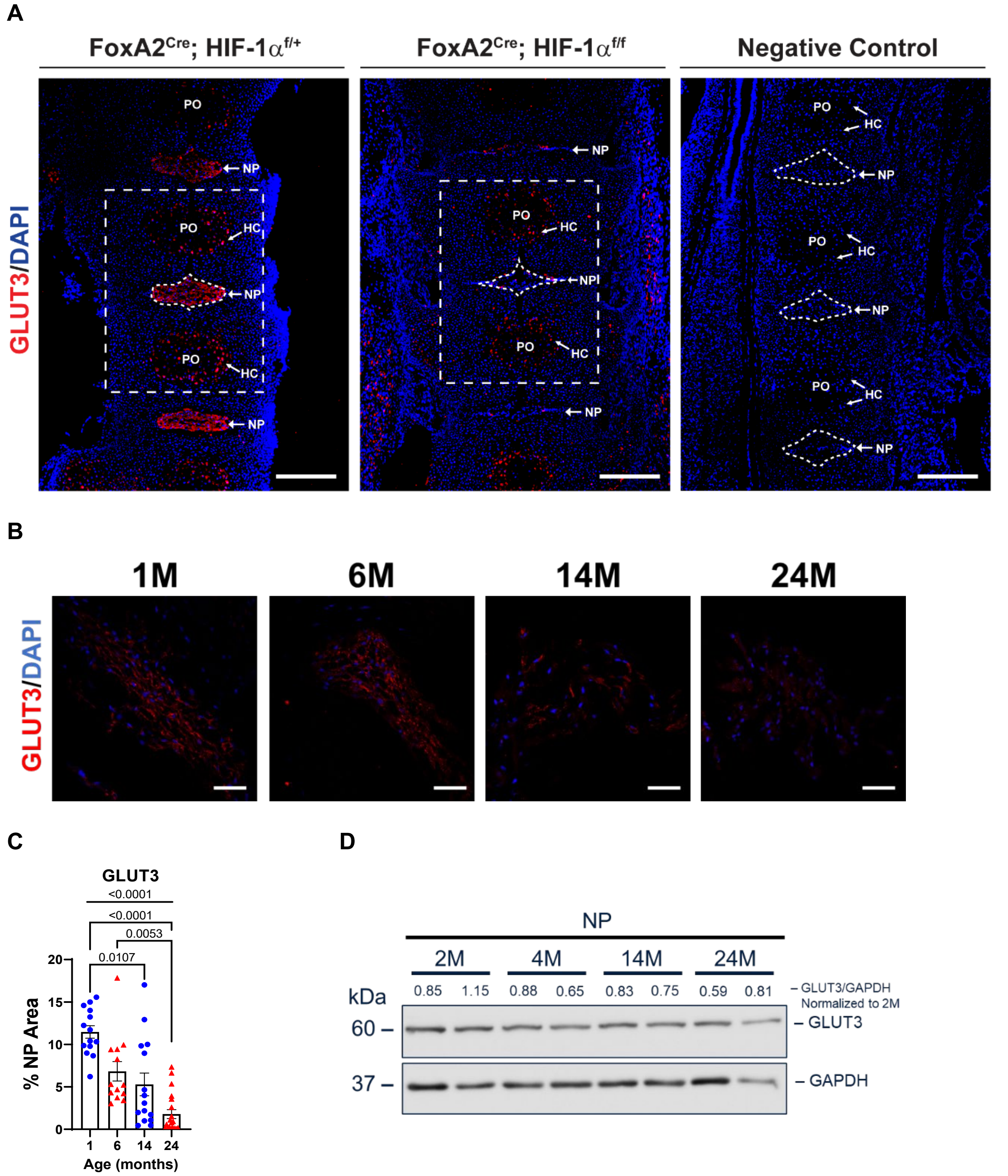


**Fig. S1. Conditional knockout of HIF-1α in NP cells leads to loss of GLUT3, and GLUT3 expression is lost with age.** (A) Example immunohistochemistry images showing glucose transporter 3 (GLUT3) expression in embryonic day 15.5 (E15.5) nucleus pulposus (NP) of FoxA2$^{Cre}$; HIF-1α$^{f/+}$ and FoxA2$^{Cre}$; HIF-1α$^{f/f}$ mice with negative control. PO, Primary center of ossification; HC, Hypertrophic Chondrocytes; NP, Nucleus Pulposus. Scale Bar = 100 μm. (B, C) Representative images and quantification of GLUT3 in wildtype (BL6/J) mice with aging (n = 5 mice/timepoint; 2-4 discs/animal, 14-20 discs/timepoint). Scale Bar = 50 μm. (D) Western blot of GLUT3 expression in wildtype (BL6/J) mice with aging, from 2-months (2M) to 24-months (24M) (n = 2 mice/timepoint; 20 discs/animal).

**A**

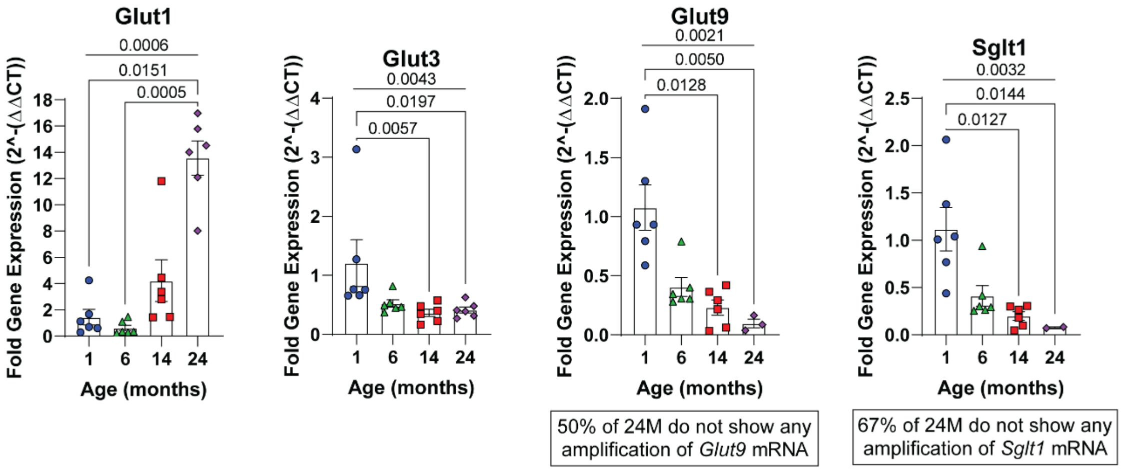


**Fig. S2. mRNA expression of glucose transporters in wildtype (BL6/J) mice with aging.** (A) Quantitative real time polymerase chain reaction (qRT-PCR) of glucose transporters, glucose transporter 1 (*Glut1*), glucose transporter 3 (*Glut3*), glucose transporter 9 (*Glut9*), and sodium/glucose cotransporter 1 (*Slgt1*) in 1-month (1M), 6-month (6M), 14-month (14M), and 24-month (24M) wildtype (BL6/J) mice (n = 6 mice/genotype; 20 disc/animal).

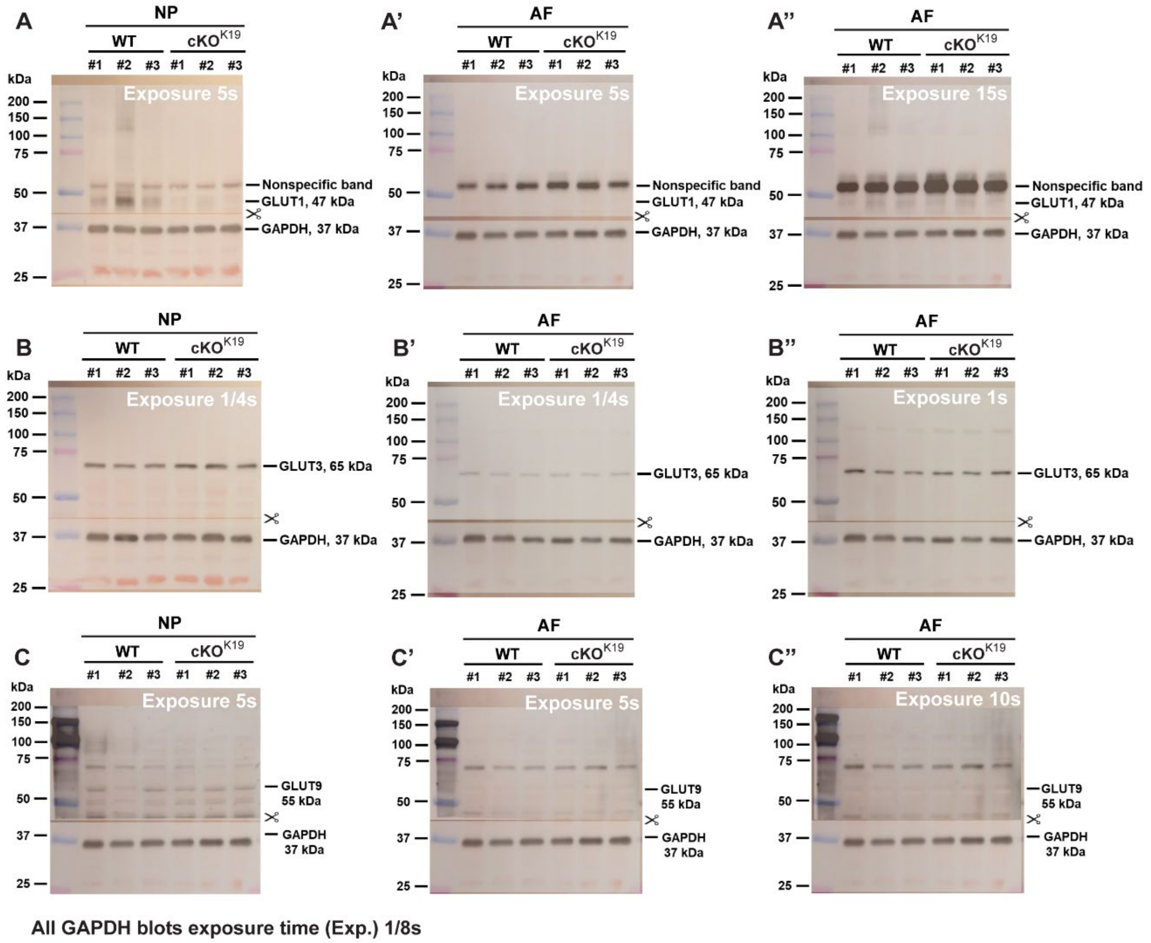


**Fig. S3. Western Blots showing GLUT1, 3 and 9 levels in NP and AF of WT and Glut1cKO$^{K19}$.** Glucose transporter 1 (GLUT1) levels in wildtype (WT, Glut1$^{f/f}$) and Keratin-19-Cre-Glut1-conditional knockout (Glut1cKO$^{K19}$) (A) nucleus pulposus (NP) and (A', A'') annulus fibrosus (AF). GLUT3 levels in Glut1 WT and Glut1cKO$^{K19}$ (B) NP and (B', B'') AF. Glucose transporter 9 (GLUT9) levels in GLUT1 WT and Glut1cKO$^{K19}$ (C) NP and (C', C'') AF. AF and NP blots are run with identical amount of tissue protein. AF blot shows identical exposure time of 5 seconds and a higher exposure time of 15 seconds to that of NP blot. All glyceraldehyde 3-phosphate dehydrogenase (GAPDH) blots had identical exposure time of 1/8 seconds. All blots are overlaid with images of Ponceau Red stained membranes to show molecular weight markers and indicates position where membranes were cut. Blots shown are for n = 3 mice/genotype; 20 discs/animal.

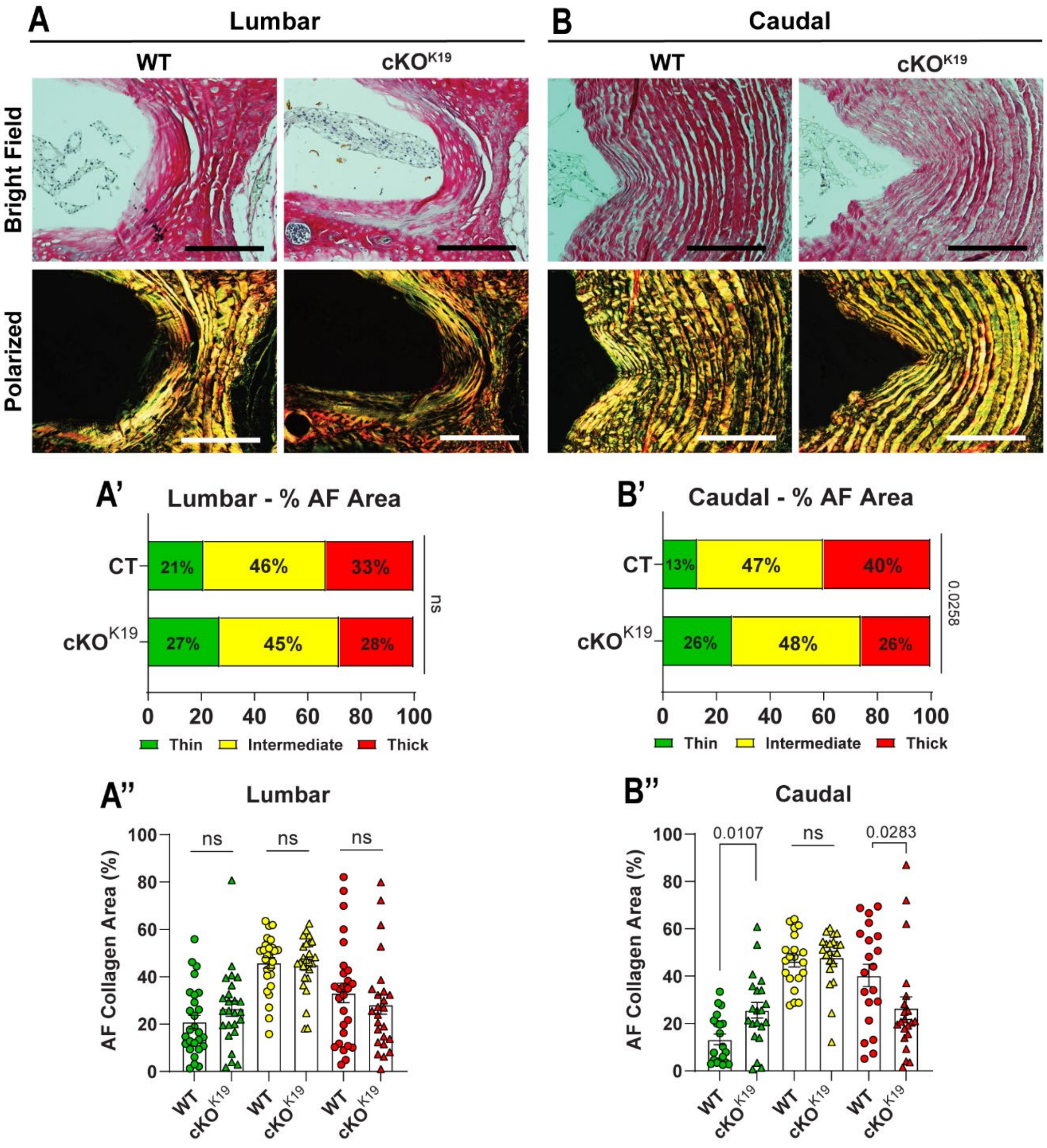


**Fig. S4. Picrosirius Red staining and polarized light imaging of Glut1cKO$^{K19}$ discs.** (A, B) Representative brightfield and polarized light images of Glut1 wildtype (WT) and Keratin-19-Cre-Glut1-conditional knockout (Glut1cKO$^{K19}$) lumbar and caudal discs. (A', A'', B', B'') Quantification of distribution and individual abundance of thin, intermediate, and thick fibers. (n = 8 WT, 7 Glut1cKO$^{K19}$ mice; 6 lumbar and 3 caudal discs/animal). Statistical significance in distribution was determined using a $\chi^2$ test. Statistical significance of thin, intermediate, and thick fiber size abundance was determined using Mann-Whitney test. Quantitative measurements represent mean ± SEM.

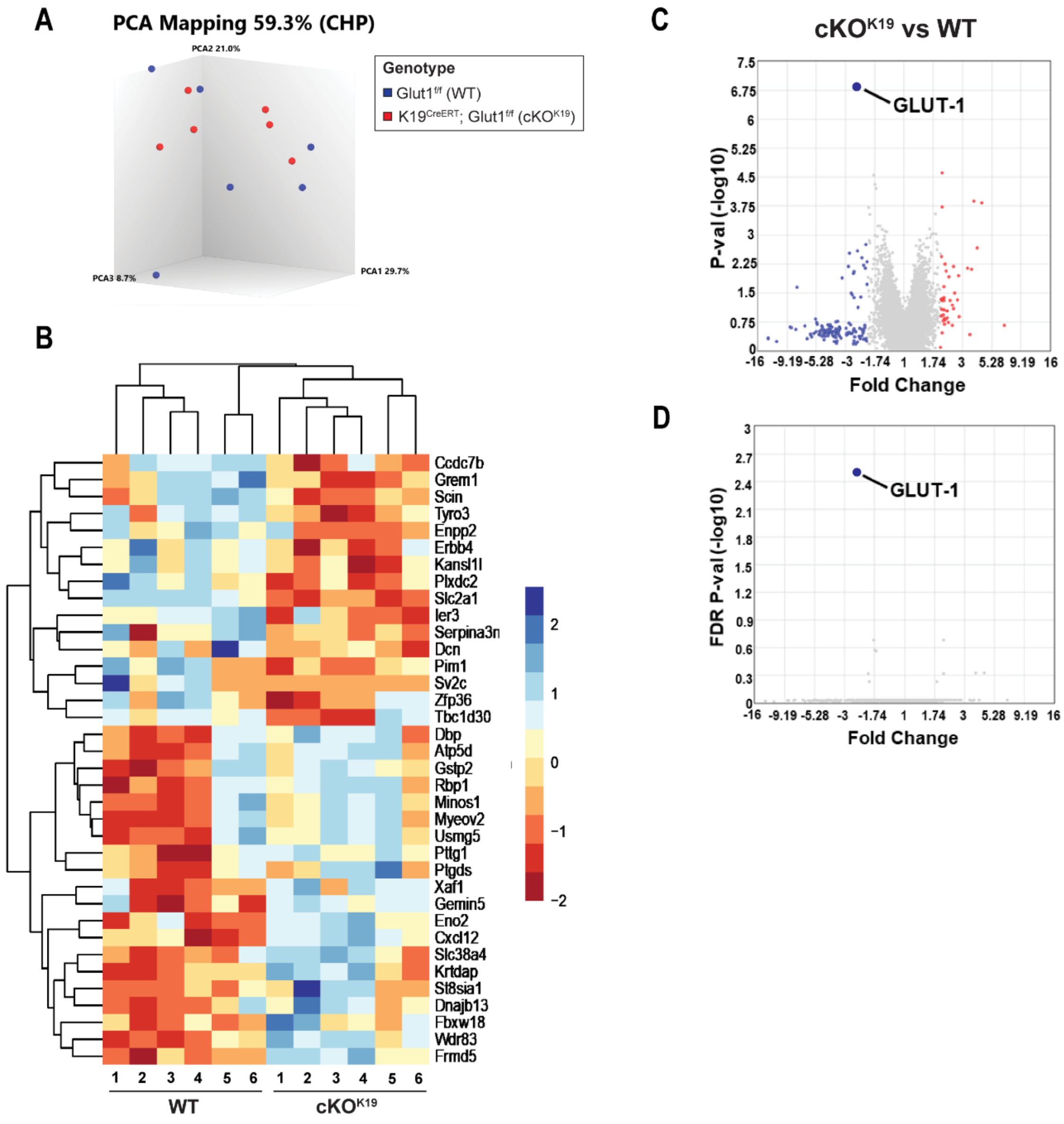


**Fig. S5. Glut1cKO$^{K19}$ mice have a similar transcriptomic profile to that of Wildtype mice.** (A) Transcriptomic profiles from 6 wildtype (WT) and 6 Keratin-19-Cre-Glut1-conditional knockout (Glut1cKO$^{K19}$, cKO $^{K19}$) mice were compared by three-dimensional principal component analysis (PCA). (B) Heatmaps of differentially expressed genes (DEGs) using Z-score between Wildtype and Glut1cKO$^{K19}$. (C) Volcano Plot showing the relationship between fold change ($\geq$ ±2-fold) and p-value (< 0.05), Glucose transporter 1 (Glut1/*Slc2a1*) is pointed out along with significantly upregulated (red dots) and downregulated (blue dots) DEGs. (D) Volcano Plot showing the relationship between fold change (>±2-fold) and false discovery rate (FDR) p-value (<0.05), the only gene meeting this FDR significance cutoff is Glut1/*Slc2a1*. (n = 6 mice/genotype; 20 discs/animal).

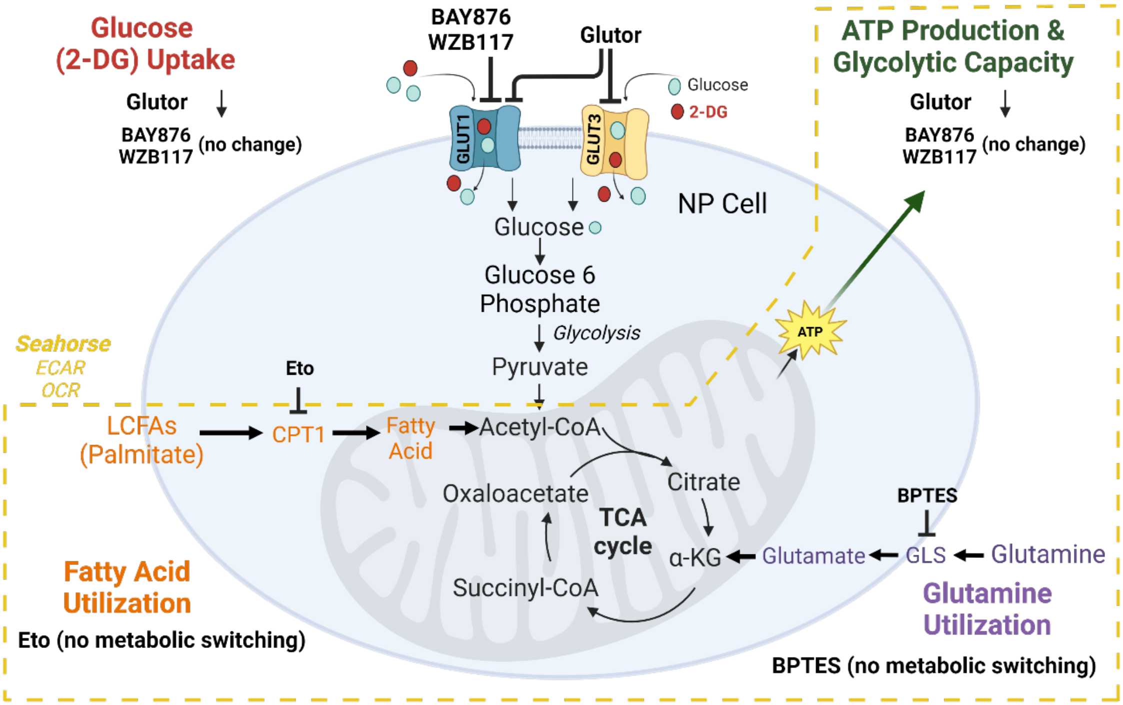


**Fig. S6. Schematic summary of the in vitro approaches and the glycolysis flux with the different substrates/metabolites measured and their outcomes.** Created with BioRender.com

**Table S1. qPCR Primers**

| Gene | Forward (5' to 3') | Reverse (5' to 3') |
|---|---|---|
| *Glut1* | GGCCTGACTACTGGCTTTGT | TGCATTGCCCATGATGGAGT |
| *Glut3* | GGTGGAGCGGTGAAGATCAG | GAGATGGGGTCACCTTCGTT |
| *Glut9* | GATGATGTCTGTCCTGGATGTAG | GAGTGATGTCGGGTGTCTTT |
| *Sglt1* | GGATCAGGTCATTGTGCAGC | TGGTGTGCCGCAGTATTTCT |
| *Hprt* | CGAGATGTCATGAAGGAGATGG | AGCAGGTCAGCAAAGAACTTA |
